# Architecture of the Neuro-Glia-Vascular System

**DOI:** 10.1101/2021.01.19.427241

**Authors:** Eleftherios Zisis, Daniel Keller, Lida Kanari, Alexis Arnaudon, Michael Gevaert, Thomas Delemontex, Benoît Coste, Alessandro Foni, Marwan Abdellah, Corrado Calì, Kathryn Hess, Pierre Julius Magistretti, Felix Schürmann, Henry Markram

## Abstract

Astrocytes connect the vasculature to neurons and mediate the supply of nutrients and biochemicals. They also remove metabolites from the neurons and extracellular environment. They are involved in a growing number of physiological and pathophysiological processes. Understanding the biophysical, physiological, and molecular interactions in this neuro-glia-vascular ensemble (NGV) and how they support brain function is severely restricted by the lack of detailed cytoarchitecture. To address this problem, we used data from multiple sources to create a data-driven digital reconstruction of the NGV at micrometer anatomical resolution. We reconstructed 0.2 mm^3^ of rat somatosensory cortical tissue with approximately 16000 morphologically detailed neurons, its microvasculature, and approximately 2500 morphologically detailed protoplasmic astrocytes. The consistency of the reconstruction with a wide array of experimental measurements allows novel predictions of the numbers and locations of astrocytes and astrocytic processes that support different types of neurons. This allows anatomical reconstruction of the spatial microdomains of astrocytes and their overlapping regions. The number and locations of end-feet connecting each astrocyte to the vasculature can be determined as well as the extent to which they cover the microvasculature. The structural analysis of the NGV circuit showed that astrocytic shape and numbers are constrained by vasculature’s spatial occupancy and their functional role to form NGV connections. The digital reconstruction of the NGV is a resource that will enable a better understanding of the anatomical principles and geometric constraints which govern how astrocytes support brain function.

**Table of contents:** *Main points:*

- The Blue Brain Project digitally reconstructs a part of neocortical Neuro-Glia-Vascular organization
- Interdependencies and topological methods allow dense in silico reconstruction from sparse experimental data
- The polarized role of protoplasmic astrocytes constrains their shapes and numbers

*Table of contents image:* 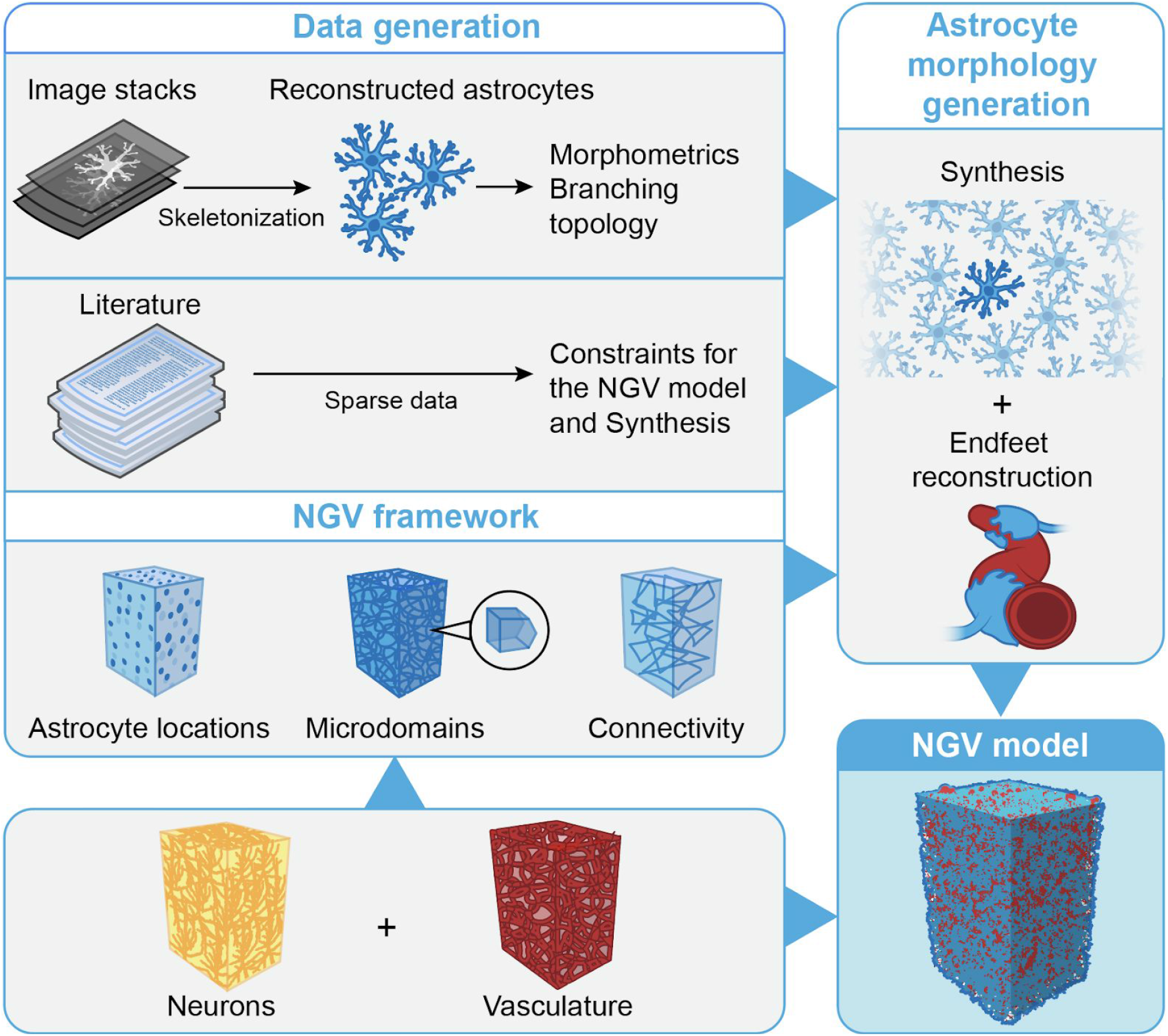

## Introduction

Research on brain function commonly focuses on just one of its cell components, the neuron, treating it as a standalone and non-interactive unit. However, this approach is prone to miss critical interactions that may affect function, such as those between cell types. Neurons, the principal subjects of neuroscience, are structurally and functionally linked to glia and the micro-vasculature, forming a complex system of multi-directional communication known as the neuro-glia-vascular (NGV) ensemble (Verkhratsky & Toescu, 2006). Studying this system requires the combination of electrophysiological and high-resolution imaging techniques to monitor the NGV physiology, a technically challenging and expensive endeavor that has started to be tackled in recent decades (Zonta et al., 2003). An anatomically accurate model of all the structural components is needed to understand and investigate normal brain mechanisms and pathologies linked to the NGV. This work provides a data-driven, biologically realistic model of NGV structural organization, which includes the three major components of the gray matter of P14 rats: neurons, astrocytes, and microvasculature.

Over the past decade, research has challenged earlier dogma and shown the nature of glia to be as intricate as that of their well-studied neighbors, the neurons (Volterra & Meldolesi, 2005). One type of glial cell, the astrocyte, links neuronal synapses to the cerebrovascular basement membrane (Figure 1A). Astrocytes are classified into fibrous and protoplasmic, based on their morphological phenotype and anatomical location (Privat & Rataboul, 2012). Fibrous astrocytes extend straight processes and are mainly located in the white matter, whereas protoplasmic astrocytes possess spongiform processes and are mainly located in the gray matter.

**Figure 1:**
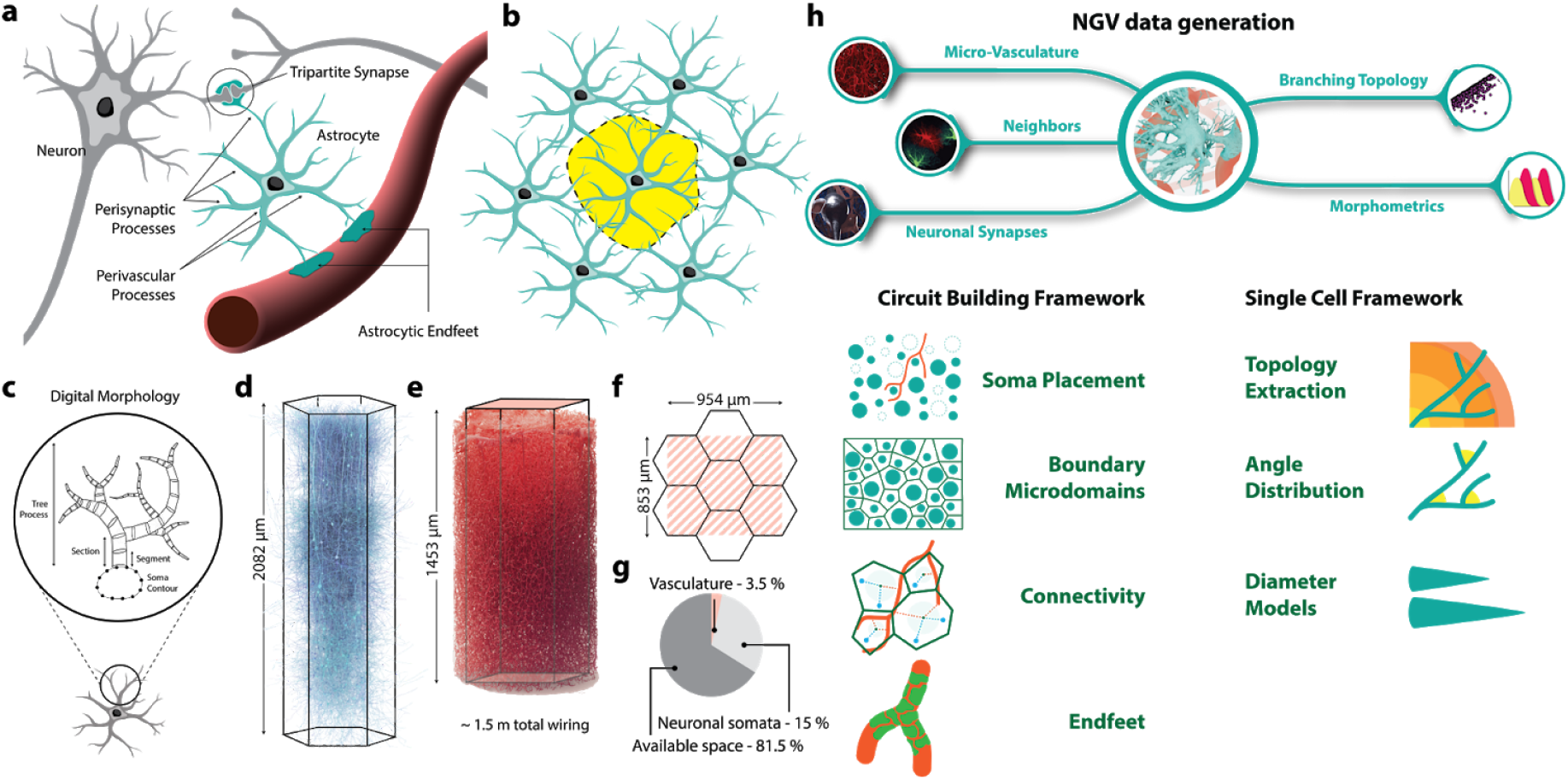
Neuro - Glia - Vascular architecture overview (a) Astrocytes contact and wrap around synapses and project their perivascular processes to the surface of the vasculature, where they form endfeet. (b) Astrocytes establish anatomically exclusive domains, which minimally overlap with their astrocyte neighbors. (c) In-silico or digital morphologies are tree structures that connect to a central soma geometry. (d) Neuronal microcircuit (e) Cerebral microvasculature (f) Overlap of neuronal mesocircuit and vasculature dataset (g) Percentage of existing volume occupancy in the circuit space (h) Building astrocytic networks requires data that is classified into two parts: Circuit building data and single-cell morphology data.

Protoplasmic astrocytes radially extend between five and ten primary processes (Calì et al., 2019; Damoiseaux & Greicius, 2009, 2009; Di Benedetto et al., 2016; Moye et al., 2019), which ramify progressively into finer and finer branches, filling up their entire spatial extent (Bushong et al., 2002). Their fine leaflet-like processes wrap around the axon-spine interface (Genoud et al., 2006; Ventura & Harris, 1999) forming tripartite units (Araque et al., 1999; Santello et al., 2012). They also extend one to five thick processes to capillaries, arterioles, and venules, attaching firmly on their surface (Abbott et al., 2006; Kacem et al., 1998; Mathiisen et al., 2010). Each astrocyte is in contact with six to fifteen astrocyte neighbors (Xu et al., 2010), forming a network that is known as the astrocytic syncytium (Figure 1B).

Although astrocytes are a vastly heterogeneous population derived from embryonic (Ge et al., 2012; Magavi et al., 2012) and postnatal progenitors (Clavreul et al., 2019; Gressens et al., 1992; Levison & Goldman, 1993), their morphogenesis mainly depends on interactions with the surrounding environment and especially with their astrocytic neighbors. More specifically, they tile the entire gray matter, forming uninterrupted regions that are mostly anatomically exclusive (Bushong et al., 2002, 2004), known as microdomains. They achieve this via a repulsion process known as contact spacing (Distler et al., 1991; Tout et al., 1993), resulting in tiling shapes that are macroscopically unaffected by nearby neuronal structures (Chang Ling & Stone, 1991; Tout et al., 1993). A thin overlapping interface is formed in their periphery where they adhere to their homotypic neighbors, establishing sub-cellular highways via process-to-process gap junctions (Nagy & Rash, 2000). This overlapping patch covers approximately 5% of the astrocyte’s territory and has been observed across brain regions and species (Bushong et al., 2002, 2004; Halassa, Fellin, & Haydon, 2007; Hara et al., 2017; Khakh & Sofroniew, 2015; Nimmerjahn et al., 2004; Sofroniew & Vinters, 2010). The formation of microdomains leads to an organized spacing among the astroglial somata, which varies with laminar depth in the cortex (Lanjakornsiripan et al., 2018; López-Hidalgo et al., 2016). This morphological complexity allows for a plethora of functions ranging from trophic support and bi-directional energy modulation to synaptic transmission (Pascual et al., 2005) and plasticity (Perez-Alvarez et al., 2014) mediated by the astrocytic syncytium.

Studies have identified molecular mechanisms that drive endfeet growth (Goldman & Chiu, 1984) and by extension the maturation of the blood-brain-barrier (BBB). During development, radial glial cells (RGCs) are generated from neuroepithelial cells of the embryonic neural tube. They extend two unbranching processes, one to the ventricular wall and the other one to the pia. RGCs attach endfeet to the vasculature early on and they may retain the pial connections after differentiation to mature astroglia (Molofsky et al., 2012; Zerlin & Goldman, 1997). The number of endfeet increases postnatally with age (Nimmerjahn et al., 2004) and covers 60% of the vasculature surface (Korogod et al., 2015) in P14 rats.

Computational neuroscience has progressed since the creation of the first data-driven biological neuronal model in the 1950s by Hodgkin and Huxley (Hodgkin A. L. & Huxley A. F., 1952) and the subsequent introduction of multi-compartment modeling of digitally-reconstructed neuronal morphologies. A digital reconstruction of a morphology is a 3D trace, made either manually or using automated skeletonization techniques (Donohue & Ascoli, 2011; Halavi et al., 2012). The branching morphology is represented as a sequence of segments (truncated cones) that capture the X, Y, Z coordinates, cross-sectional diameter, and connectivity links (Figure 1C). Biologically realistic neuronal morphologies allow for compartmental biophysical models, which can capture the non-linear dynamics arising from the distribution of ion channels throughout their compartments. The branching structure of neuronal morphologies is believed to play an important role in the computational tasks of the neuronal networks (Cuntz et al., 2007; van Elburg & van Ooyen, 2010). By the same rationale, the branching topology and geometry of astrocytic morphologies are important for driving the propagation of calcium-induced waves that have been found to contribute to information processing (Araque et al., 2014; Cornell-Bell et al., 1990; Hamilton & Attwell, 2010; Volterra & Meldolesi, 2005), as well as biochemical compartmentalizations such as those involved in energy metabolism (Magistretti & Allaman, 2018). However, because of the densely ramified nature of astrocytes, digital reconstructions using markers such as GFAP only capture the primary processes of the actual morphologies (Kulkarni et al., 2015). Three-dimensional electron microscopy on the other hand can provide the ultrastructure required for capturing the nanoscopic astrocytic processes but at the cost of low throughput (Calì et al., 2019).

In this work, we generated for the first time an *in-silico* anatomical reconstruction of the P14 rat gray matter, comprised of neurons, protoplasmic astrocytes, and the cerebral microvasculature at the level of biophysically-detailed morphologies. To address the intricate structural organization of the NGV components, we combined prior work on the reconstruction of neuronal microcircuitry (Markram et al., 2015) with sparse literature data to develop a large-scale algorithmic framework for growing detailed astrocytic morphologies that connect to neurons, the vasculature, and other astrocytes while tiling the cortical space.

To achieve this we first reconstructed the population-wide organization of the NGV. We co-localized neuronal and the vascular datasets (Figure 1e-g) in a shared region, which was then populated with astrocytic somata. Next, we partitioned the cortical space into tiling and overlapping polygons, representing the microdomains of astrocytes. Within the boundary of each domain, the connectivity of the somata to synapses and to the vasculature was established.

To grow detailed astrocytic morphologies we first solved the challenge of the small number of experimental astroglial morphologies, by using topology to reproduce the branching pattern of the astrocytes (Kanari et al., 2018). We combined this approach with the population-wide data and geometrical constraints generated by the steps above into a novel synthesis algorithm that allowed to *in-silico* grow astrocytic morphologies in their local environment (Figure 1h). They formed connections with neuronal synapses, projected processes to the vasculature, and grew endfeet that wrapped around the surface of the vessels. We aimed to generate morphologically accurate astroglial cells *in-silico*, rather than model their development.

We demonstrated the predictive power of the NGV circuit by investigating the compositional and organizational principles that govern the underlying biological complexity. We analyzed the co-localization of astrocytic somata, large vessels, capillaries, and endfeet targets on the vasculature’s surface in search of the dominating element in the endfeet organization. We extracted the total lengths, surface areas, and volumes of the segments of neurons, astrocytes, and the vasculature to explore the composition of geometrical features in the neocortical space and their relationship. Focusing around the central player of the NGV, the astrocyte, we extracted connectivity statistics, opening a window to the NGV network organization. Finally, the NGV model exploration was extended to the generation of multiple circuits of increasing densities to investigate the astrocyte numbers effect on the endfeet connectivity and microdomain packing.

A large-scale anatomical reconstruction of such a complex organization has not been possible before; however, it is an essential step towards understanding how our brain works in health and disease. Most importantly, the NGV framework enables the simulation of physiology throughout an entire region, embedded in its actual cortical space with a biologically realistic spatial architecture, and offers the possibility to shed light on unknown questions about the microscopic brain interactions.

## Materials and Methods

### NGV input datasets

#### Voxelized virtual brain atlas equipped with 3D density profiles

For the construction of circuits, a 3D voxel atlas was generated covering the volume of interest, which spanned a region of 954μm × 1453μm × 853μm. This bounding region was determined by co-localizing the vasculature dataset (Reichold et al., 2009) with a neuronal mesocircuit (Markram et al., 2015), which consisted of the central hexagonal microcircuit surrounded by six satellite ones. Both datasets were aligned to the pia and the maximum region was determined so that it includes both vasculature wiring and neuronal somata without empty locations.

The x, z axes of the region’s coordinate system were parallel to the pial surface, whereas the y axis was perpendicular to it. The voxel dimensions of the atlas were chosen to 10μm × 5μm × 10μm, where the voxel y dimension (depth) was determined so that it would reflect the discretization step of the density profile that was used as described below.

One-dimensional astrocytic densities (Number of astrocytes per mm^3^) along the y-axis (Appaix et al., 2012) were mapped to the 3D voxelized grid. Voxels lying on the xz plane, parallel to the pia, were assigned a constant density, while voxels along the y-direction matched the density from the input profile.

#### Digital microvascular network skeleton and surface meshing

The cerebral microvasculature plays a vital role in the structural topology of NGV networks during the developmental and maturation stages of the CNS. In our models, we distinguish between two types of datasets: the vascular skeleton and the surface mesh. The vasculature skeleton is represented as a graph of consecutive point chains (Figure 2A-C), which is oriented and can contain cycles. Each point is also associated with a diameter, which reflects the cross-sectional width of the vessel at that point. Although most of the steps in the circuit building pipeline use the skeletonized representation of the cerebral microvasculature, the generation of endfeet appositions on the surface of the vasculature requires a more detailed representation of the surface geometry. Thus, starting from the skeletonized dataset we reconstructed a triangular discretization of the surface geometry with variable resolution. The surface mesh was generated based on implicit structures, known as metaobjects (Abdellah et al., 2020; Oeltze & Preim, 2004), which allowed the creation of highly-detailed meshes of vasculature datasets (Figure 2D-F).

**Figure 2:**
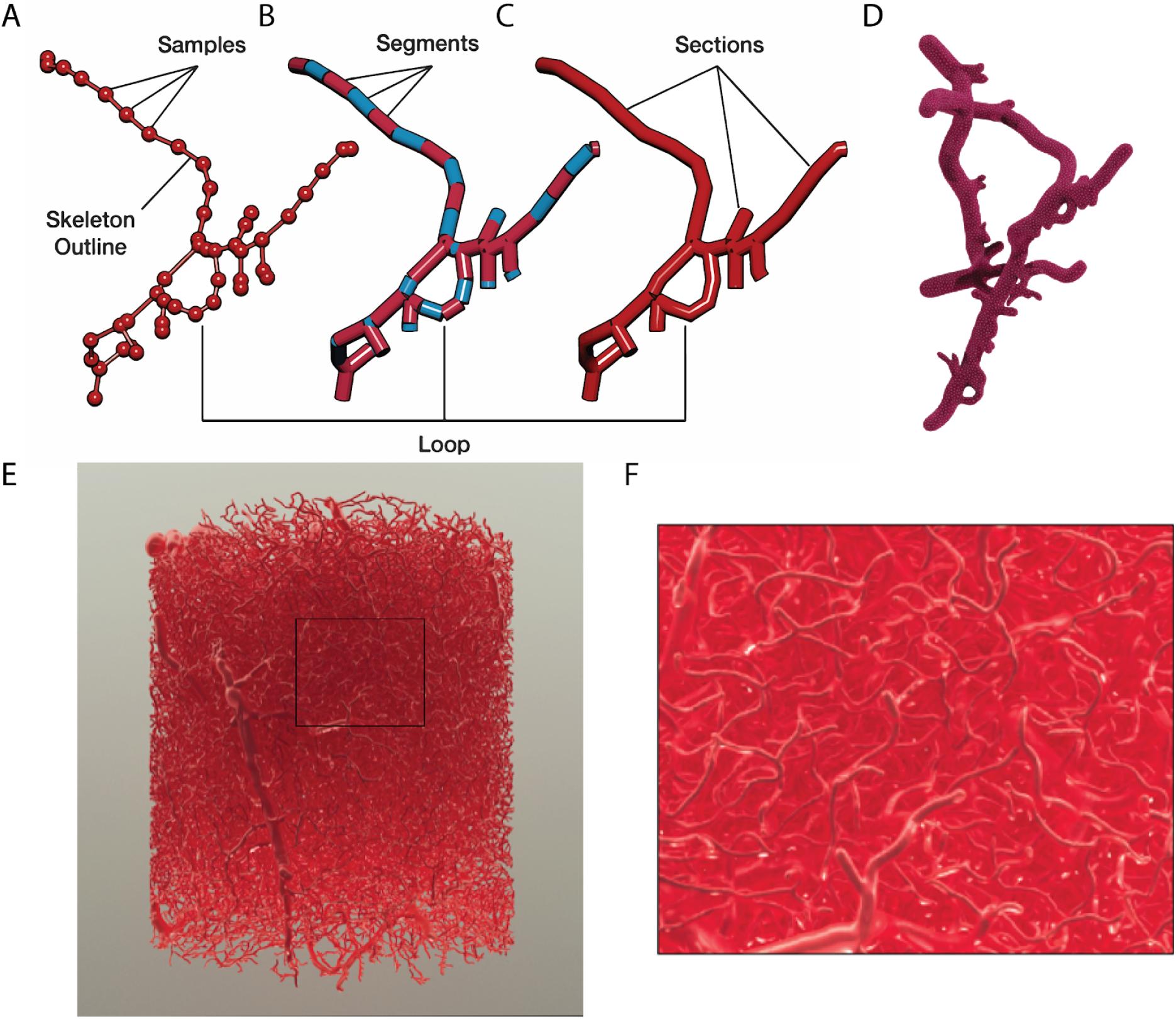
Vasculature meshing (A) The vasculature dataset consists of points with a diameter linked by edges. (B) Two consecutive points linked by an edge are defined as a segment. (C) A chain of consecutive points between two forking points is defined as a section. (D) Generated surface mesh with overlaid triangles. (E-F) Original vasculature dataset mesh.

#### Experimental morphologies of astrocytes

Image stacks of astrocytic morphologies from P14 rats were provided by the laboratory of Prof. Pierre Magistretti (King Abdullah University of Science and Technology). They were extracted using high-resolution serial block-face imaging, capturing the nanoscale structure of astrocytes with a 20 nm resolution (Calì et al., 2019).

The stacks were converted into surface meshes, and the geometry was verified to be watertight and manifold. Finally, the surface meshes were contracted into skeletons consisting of nodes (coordinates and radii) and their connectivity (Shapira et al., 2008).

The reconstructed astrocyte morphologies were used both for the extraction of the branching topology of the P14 rat astrocytes (see Branching topology analysis) and for the calculation of morphometrics used in the topological synthesis and validation.

#### Neuronal neocortical mesocircuit

Neuronal neocortical circuits of the somatosensory cortex of the juvenile rat were algorithmically generated based on the circuit building framework that was previously published by Markram et al., (2015). In this work the input neocortical mesocircuit consisted of a central microcircuit, occupying a volume of 0.28 mm^3^, with 23,590 neurons surrounded by six satellite microcircuits of 139,992 neurons in total. The overlapping arbors of neuronal morphologies formed ~8 million connections with ~37 million synapses, constituting the basis of the neuronal component of the NGV networks and providing the synaptic locations that are required from the NGV connectome.

#### Topological analysis of experimental astrocytes

In their work, Kanari et al., (2020) have shown that the distinct branching shapes of neuronal dendrites can be reproduced using the topological morphological descriptor (TMD) method (Kanari et al., 2018), which extracts a topological representation (persistence barcode) of a tree’s branching structure. A barcode is a collection of bars, with each bar encoding the start (root or bifurcation) and the end (termination) path lengths of a branch. The barcode is used to define the bifurcation and termination probabilities of each branch during the computational growth of morphologies. The coupling of these probabilities provides a method to implicitly reproduce key correlations between morphological features.

In order to capture the branching topology of astrocytes, we applied the TMD method on experimentally reconstructed astrocytes. Because of their different structural role, perivascular and perisynaptic trees (Figure 3A) were converted separately into two collections of barcodes (Figure 3B). Each collection was then used in the synthesis of new astrocyte morphologies to reproduce their branching structure.

**Figure 3.**
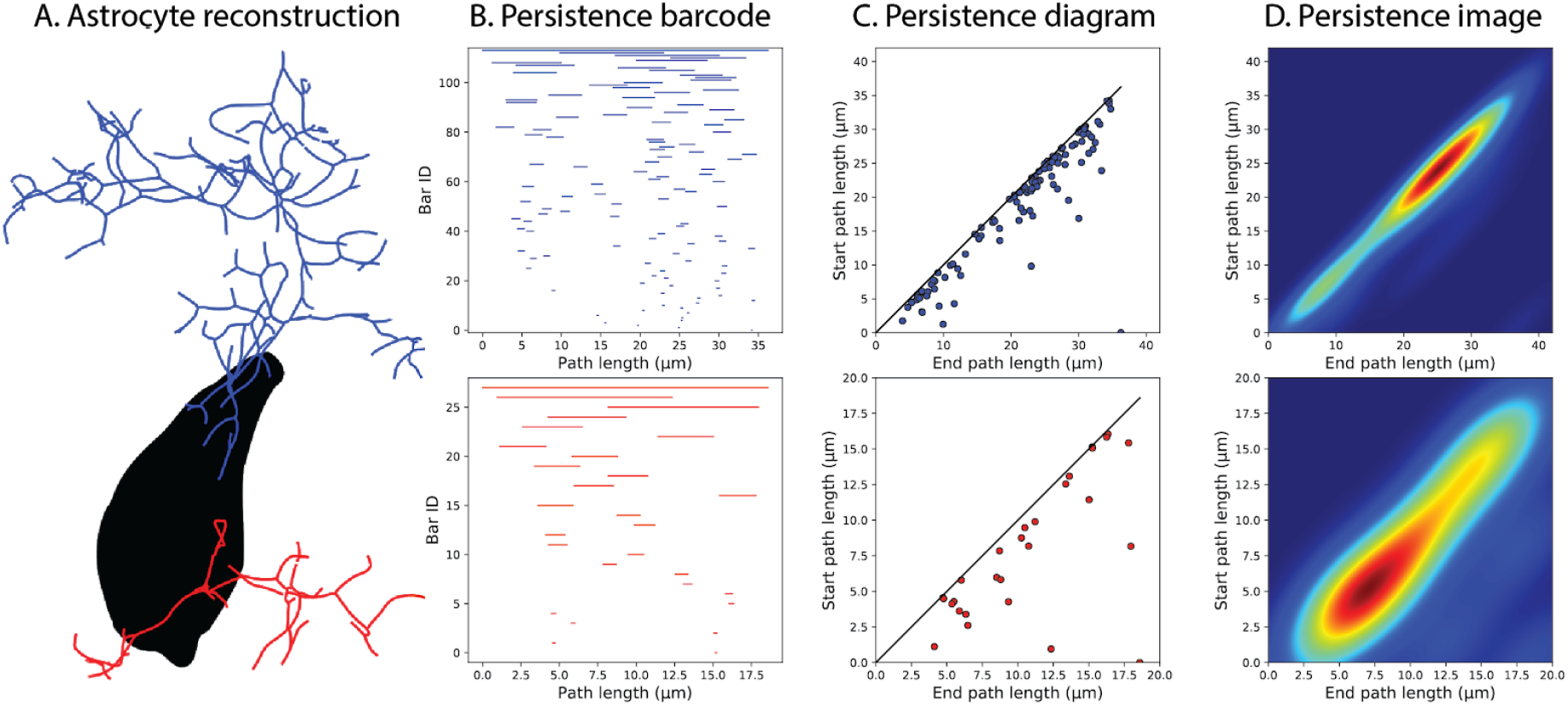
Example of the decomposition of astrocytic trees into persistence barcodes. (A) Two small trees of perivascular (blue) and perisynaptic (red) types were selected for demonstration purposes from an experimentally reconstructed astrocyte morphology (soma in black). (B) Each tree is decomposed into a barcode, with each branch being represented as a horizontal line (bar) marking the start and end path length from the soma. The barcode in (B) can also be represented as points in the persistence diagram (C), in which the start and end path lengths of each bar are shown as y and × coordinates respectively. (D) By applying a kernel density estimator on the persistence diagram the persistence image is generated, which shows the bar density of the branching structure.

Persistence barcodes are alternatively represented as persistence diagrams, in which the start and end points correspond to the y and x-axis of a 2D diagram (Figure 3C). The diagonal on the persistence diagram corresponds to bars that have equal start and end path lengths, i.e. zero length. Applying a kernel density estimator on the persistence diagram results in a persistence image (Figure 3D), which shows the bar density as a 2D pixel image. The persistence image was used in topological validation because it allows calculating a topological distance between two images from the difference of the pairwise pixel values.

### Reconstruction algorithms

#### Generating astrocytic positions without collisions

Astrocytes exhibit variation in both density (Appaix et al., 2012) and dispersion because of their tiling organization (Bushong et al., 2002, 2004). Previous placement algorithms placed cells with respect to their density (Erö et al., 2018; Markram et al., 2015) across layers and brain regions. However, astrocytic spacing results in a nearest neighbor distance distribution that is not uniform and depends on animal species and age (Distler et al., 1991; López-Hidalgo et al., 2016).

To address this, we developed a new stochastic placement method, which models astrocytic organization as a spatial random pattern *X* (Baddeley et al., 2007), the configuration p of which is governed by a Gibbs energy functional *E(p)* (Dereudre, 2019; Ruelle, 1999):

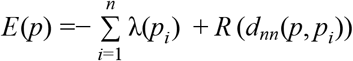

where λ(*p*_*i*_) is the astrocytic density, *d*_*nn*_(*p*, *p*_*i*_) is the distance of the location *p*_*i*_ to its closest neighbor, and *R* is a repulsion potential. The probability *P* (*X* = *p*) of the random pattern being in this configuration is given as:

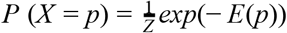

where *Z* the normalization constant known as the partition function, which sums all possible configurations of the pattern. Because of the dependence of the configuration energy from both density and pairwise repulsion, direct sampling from the target distribution was not possible. Therefore, we sampled somata positions from the Gibbs distribution using the Metropolis-Hastings algorithm (Robert & Casella, 1999).

Starting from a voxelized region equipped with a spatial density profile, we calculate the total number of astrocytes per voxel (Figure 4A). Each time a soma is placed, the respective voxel count is reduced by one (Figure 4A-B). For each placement trial, a position is chosen first by randomly selecting a non-occupied voxel and then by uniformly sampling a point inside that voxel. A radius is also sampled from a normal distribution with a mean of 5.6μm and a standard deviation of 0.7μm. A collision of the newly generated soma with the vasculature geometry or already placed astrocytic somata would result in its rejection and the process would repeat until a trial soma is found that doesn’t collide with other geometries.

**Figure 4:**
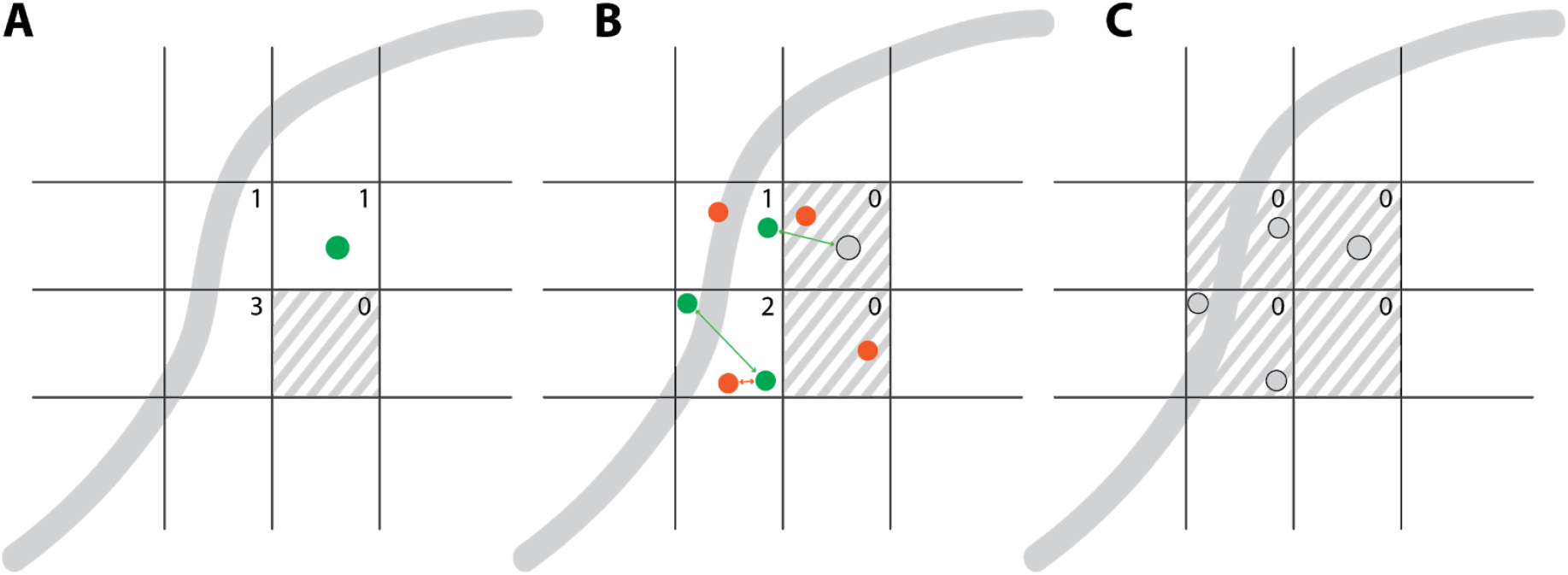
Astrocytic somata placement procedure. Squares represent the voxels of the atlas and the numbers are the expected astrocyte counts as calculated from the astrocytic density. Green circles represent accepted soma placement trials, red the rejected, and gray the existing ones at the current step. Similarly, green and red lines represent accepted and rejected nearest distance evaluations respectively. The continuous gray line represents the geometry of a vessel. (A) The astrocytic density is converted into astrocyte counts, which are decremented when a soma is placed into the respective voxel. (B) A placement trial is rejected if it is placed into a maxed-out voxel, or if it collides with other geometrical entities. (C) Placement stops when either all the voxels have been filled or if there is no available space to add more somata.

If there is no collision the energy functional *E(p)* is calculated. The ratio between the new over the previous energy determines the acceptance of the new trial according to the Metropolis-Hastings algorithm. If not accepted, a new position/radius combination is generated and the procedure repeats until all voxels have reached their total counts or if there is no available space left (Figure 4C).

#### Microdomain boundary for growing astrocytes

The anatomically exclusive territories of protoplasmic astrocytes arise from the physical extent of their morphologies (Bushong et al., 2002, 2004; Distler et al., 1991), tiling the entire cortical space. We modelled the microdomains as a partition of the 3D space into polygonal regions (Laguerre tessellation; Aurenhammer 1987), the geometry of which was calculated from the somata positions and radii. Contrary to a regular Voronoi tessellation, the size of convex domains in a Laguerre tessellation is proportional to the radius of the somata.

This geometrical abstraction generated tiling convex polygons, allowing for the establishment of the bounding region for each astrocyte, which was subsequently grown in it. Starting from a tiling organization of polygons and growing astrocytes in them correctly simulated the morphological organization of real astrocytes.

Furthermore, two microdomain datasets were generated: a regular and an overlapping tessellation. Overlapping microdomains were created by uniformly scaling the domains of the regular tessellation until the desired percentage of overlap is achieved. For P14 rats this overlap is approximately 5% of the domain volume (Ogata & Kosaka, 2002).

### Reconstructing the NGV connectome

Three types of connectivities were reconstructed via the NGV circuit building pipeline: gliovascular connections (astrocyte-vasculature endfoot), neuroglial connections (astrocyte-neuron tripartite synapse), and glial connections (astrocyte-astrocyte gap junctions). For their establishment, the microdomain tessellation determined the available bounding region of each astrocyte. Therefore, the vasculature sites and the synapses each astrocyte can project to, are limited to the inside of its microdomain geometry.

To establish the connectivity between astrocytes and the vasculature, we first distributed potential targets on the vasculature skeleton graph with a frequency of 0.17 μm^−1^ (McCaslin et al., 2011) and then determined which fraction of the resulting point cloud was included in each microdomain boundary polygon. Based on literature data, each astrocyte was assigned a number of endfeet, ranging from 1 to 5 (Moye et al., 2019) and the endfeet sites were sequentially selected according to the following observations and experimental astrocyte reconstructions: endfeet processes minimize their distance to the vascular site (Kacem et al., 1998), maximize the distance to nearby endfeet sites and target different branches (Calì et al., 2019).

Similarly, neuroglial connectivity was determined by first finding the synapses within each microdomain geometry. Then a 60% random subset was selected to match the experimental observations (Reichenbach et al., 2010).

Finally, gap junctions between neighboring astrocytes were determined as touches between the colliding morphologies of the grown astrocytes, using the process of touch-detection as presented in (Markram et al., 2015). For this step, the full-grown astrocyte morphology was required (see Algorithmic growing of astrocytic morphologies).

#### Algorithmic generation of astrocytic morphologies

Astrocytes initially grow primary processes, which they radially extend outwards, and then ramify extensively until they fill their territories (Bushong et al., 2002, 2004; Ogata & Kosaka, 2002). Within their domains, small branches exhibit increased coverage in proximity to synapses (Genoud et al., 2006). It has been shown in culture that astrocytes extend processes towards substances that are released by synapses (Cornell-Bell et al., 1990; Hatten, 1985; Matsutani & Yamamoto, 1997). To reproduce this behavior, we used the synapses from the neuronal circuit as attraction points to influence growth.

The morphological synthesis of astrocytes combines the topological synthesis of neurons (Kanari et al., 2020) with the resource-driven growth of the space colonization algorithm of (Runions et al., 2007), adapted for stochastically grown morphologies. Consuming resources as morphologies grow guarantees the exploration of the space by the movement of the growth front into areas that have available resources, while a topology-driven branching ensures the generation of morphologies with similar branching patterns to the experimental astrocyte reconstructions.

### Initiation of processes

Astrocytic processes can be classified into perivascular or perisynaptic processes based on whether or not they connect to the vasculature. Perivascular processes project to nearby vessels, while general processes extend to form and fill the domain of the astrocyte incorporating neuronal synapses. The initial direction of perivascular processes was dictated by the contact sites that have been established upon the generation of the gliovascular connectivity. On the other hand, the direction of the perisynaptic processes was influenced by the geometry of the domain and the existing endfeet direction so that there is no overlap with the latter (Figure 5A). To achieve this, the process directions are added iteratively by maximizing the mean angle to all existing orientations. There is also a preference for large vectors, in order to follow the elongation of the domain.

**Figure 5:**
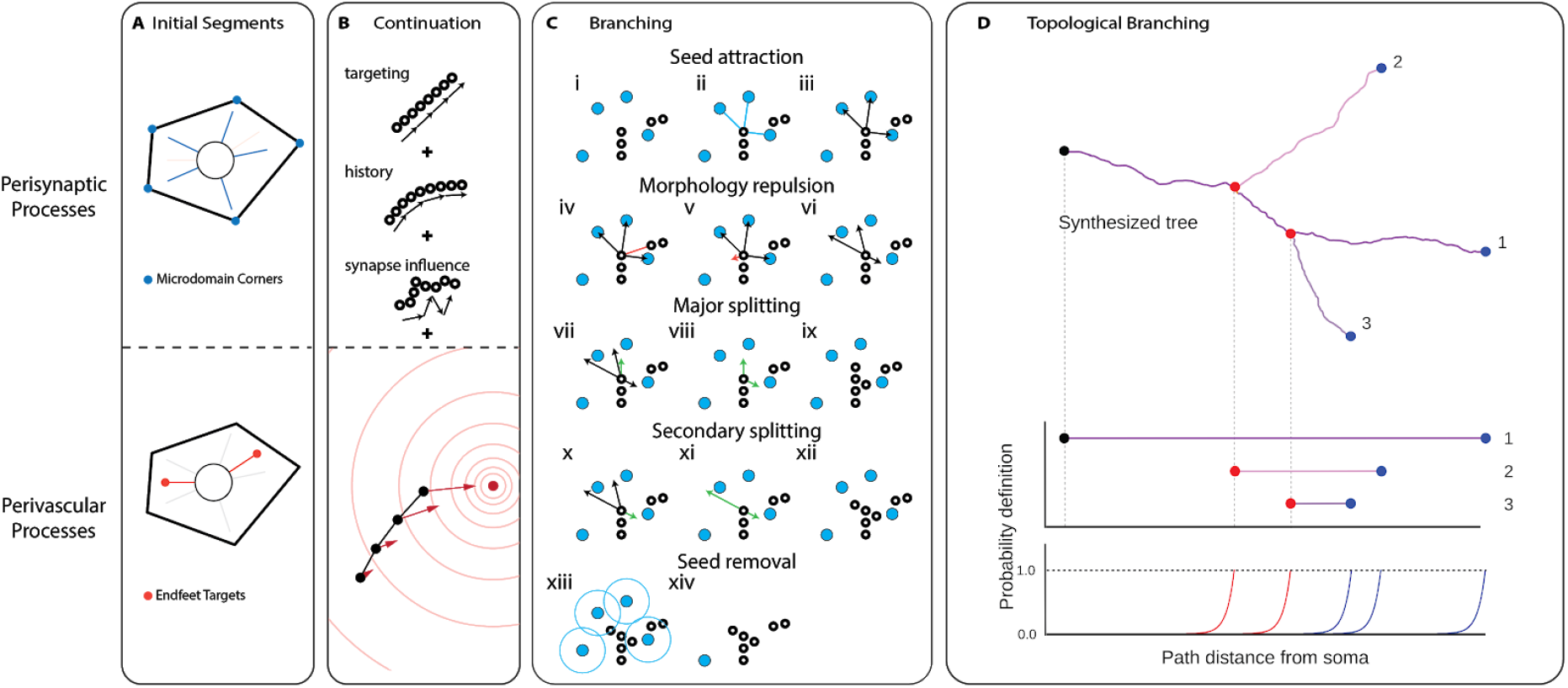
The topological exploration algorithm for synthesizing astrocytes. (A) The direction of the initial segments is chosen based on process type. Perisynaptic directions are calculated from the geometry of the domain, while perivascular ones are predetermined from the vasculature appositions. (B) The growth of processes is influenced by a targeting factor that determines the deviation to the initial orientation of the section, the history that determines the rigidity and a random influence, determined by the nearest synapses. For perivascular processes, there is an additional influence from the vasculature target that acts as an attractor to the growth. (C) To create a bifurcation, the synapses around the last point of the section are found (i-ii). The subset of resources that is within a distance of influence D_infl_ is selected (iii) creating the potential directions. (iv-vi) Each direction is influenced by a repulsion vector that is calculated from the closest morphology points within a kill distance D_kill_. The branching behavior of a section depends on its growing type, which can be either major or secondary. (vii-ix) The first direction of the major processes is always the parent direction and the second one is the most antiparallel potential seed vector. (x-xii) The first direction of the secondary processes is the direction to the closest seed and the second direction is again the most antiparallel remaining seed vector. (xiii-xiv) Following the creation of the bifurcation the synaptic points that lie within a kill radius of the morphology points are removed. (D) The branching or termination points are stochastically determined from the TMD decomposition of the digital reconstructions

### Elongation

Each process is grown point by point, forming segments. The expected segment length is drawn from a normal distribution and for astrocytes (see supplementary material for rationale), and it can be as small as 0.1 μm. The direction of the segment is a weighted sum of three unit vectors (Figure 5B): A targeting vector, the cumulative memory of the directions of previous segments within a branch and the direction to the closest synapse. Here, the randomness factor (Koene et al., 2009), which is calculated by uniformly choosing a direction on the unit sphere, has been replaced by the synaptic cloud influence.

### Bifurcation - Termination

When the branching probability, which is extracted from the barcode, triggers a bifurcation of the tip of a growing section, the splitting algorithm determines the directions and growing types of the two child sections. There are two growing types: major and secondary, which encode the behavior of the primary and higher branch order processes respectively. During initiation, all processes are given the type “major”. After a bifurcation, the types of children depend on the type of the parent section. If the latter is major, then one child is given the type major while the other one is given the secondary type. If the parent is secondary, then both children are secondary.

The first step of the branching algorithm is to find the synapses that are within an influence distance Dinfl of the process tip. The potential seed vectors are formed from the selected synapse points (Figure 5Ciii). Then, each direction is influenced by a repulsion vector, which is computed from the morphology points that are closer to the tip than a kill distance D_kill_ (Figure 5Civ-vi). The first branching direction of major parent sections is always the parent direction and the second direction is the most antiparallel potential seed vector to the first one (Figure 5Cvii-ix). The first direction of secondary parent processes is the direction to the closest seed and the second direction is again the most antiparallel remaining seed vector (Figure 5Cx-xii). Following the creation of the bifurcation the synaptic points that lie within D_kill_ of the morphology points are removed (Figure 5Cx-xii). The removal of the synapse points that are close to the morphology as the latter grows, allows the exploration of the space, based on the remaining resources at each step of the growing algorithm.

The basic principle for the tree growth can be summarized by the following steps: the tree has a probability to bifurcate and to terminate associated with the path distance from the soma (Figure 5D) During the growth process, the topological barcode, which is extracted from the reconstructed morphology population, determines these probabilities to terminate or bifurcate. Similarly to neuronal synthesis, the bifurcation/termination probabilities depend exponentially on the path distance of the growing tip from the soma. This means that when the growing tip approaches the target bifurcation or termination distances as defined from the barcode, the probability to bifurcate or terminate increases exponentially until it reaches 1 after the target distance is surpassed.

### Surface area and volume distribution

Due to a high surface-to-volume ratio in astroglial processes, it is difficult to capture their membrane geometry using a cylinder representation. For this reason, the volume and surface of segments are separately encoded in their diameters and perimeters.

The model for generating synthetic diameters for astrocytes is based on the neuronal algorithm (Kanari et al., 2020). First, distributions of the model parameters are obtained from a fit to the available reconstructed astrocytes. Then, diameters are generated using parameters sampled from these distributions. The model parameters include the sibling ratio (ratio between diameters of daughter branches), diameter power relation (to model the relative diameters between parent and daughter branches), taper rates, trunk, and terminal diameters. See for example (Ascoli et al., 2008) for more details on these parameters. We deviated from the neuronal diameter model by allowing for negative taper rates, to obtain increasing and decreasing diameters with path distances. Along the neurites, diameters are bound by the trunk and terminal diameters, sampled for each neurite, while the other parameters are sampled at each segment or bifurcation.

Regarding the distribution of perimeters on the synthesized astrocytes, we extracted the diameter-perimeter pair values from all experimental reconstructions and fit a linear regression model:

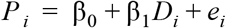

where *P*_*i*_, *D*_*i*_ are the perimeter and diameter respectively for each pair, and *e*_*i*_ is the variation or noise in the data for each node. Thus, following the diametrization process, we assigned the perimeters using the linear predictor function shown above.

### Growing endfeet surfaces

The surface geometry of the endfeet wrapping around the vasculature was generated from the positions on the surface of the vasculature (endfeet target sites, Figure 6A), which have previously been assigned in the gliovascular connectivity step. From each endfoot target site, the endfeet area is grown isotropically on the vessel surface (Figure 6B) until it collides with another endfoot area or reaches a maximum radius (Figure 6C). The growth is considered competitive because all the endfeet are growing simultaneously restricting the area they are grown into from neighboring endfeet. After the simulation has converged we prune the overshoot surfaces (Figure 6D) so that they match the experimental distribution of endfeet areas which is approximately 200 μm^2^ (Calì et al., 2019). In Figure 6E-H an actual simulation of the different color endfeet areas as they grow on the surface mesh of a test dataset can be seen.

**Figure 6:**
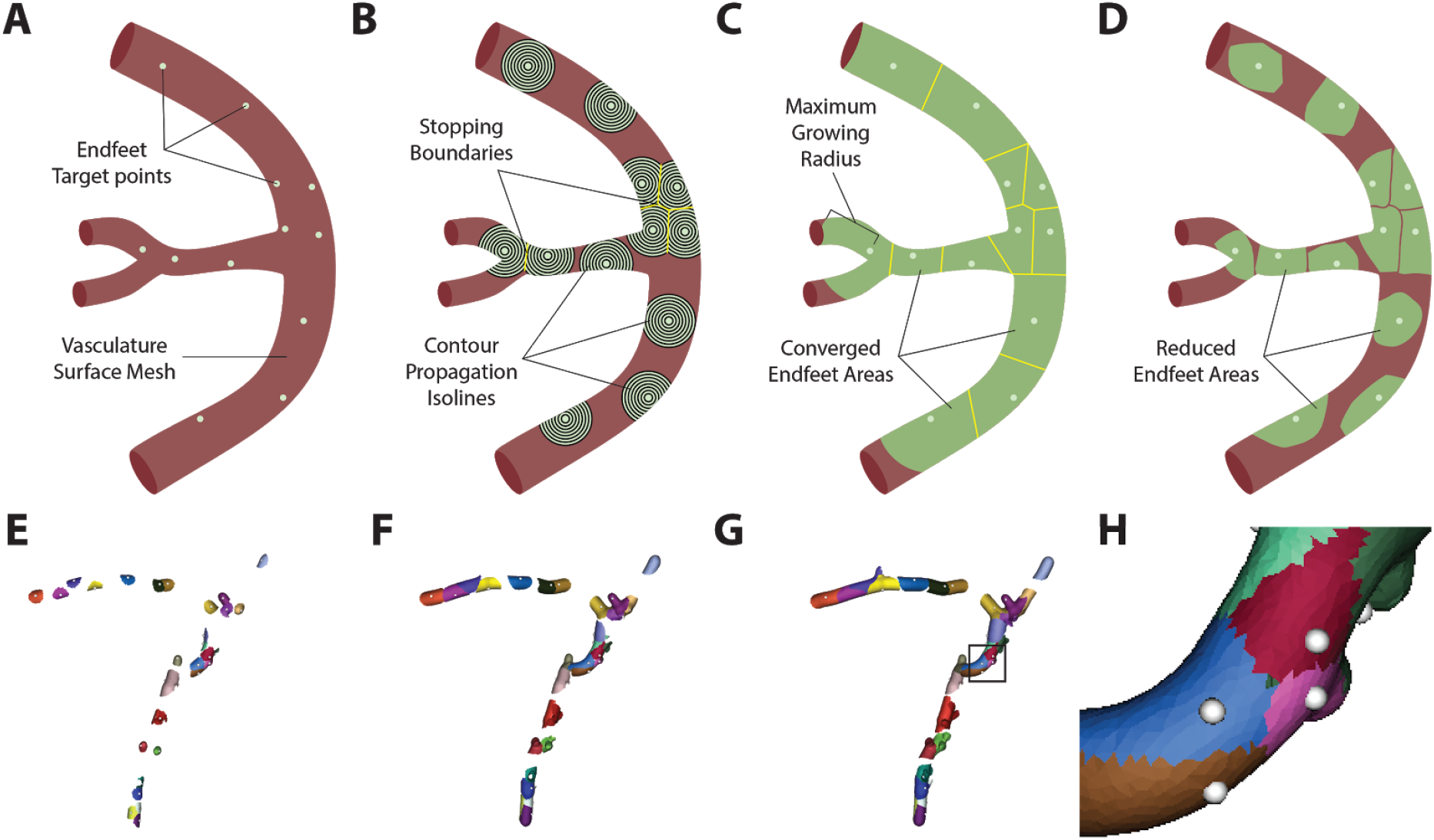
Growing of endfeet areas. Starting from the initial endfeet target points (A), waves propagate following the geodesics of the vasculature surface, until they either reach an already occupied area or exceed a maximum growing radius (C). The converged areas are then truncated so that they match the input area distribution (D). (E-H) Example of simulations with growing areas and a closeup (H) of their convergence.

#### Data availability statement

The data that support the findings of this study (experimental data, models and tools used in the circuit building process) are openly available in the NGV portal website at bbp.epfl.ch/ngv-portal.(*NGV Portal*).

## Results

The available input vasculature dataset determined the bounding space of the generated NGV circuit, the dimensions of which are 955 × 1452 × 853 μm^3^, corresponding to a volume of 1.18 mm^3^, approximately 0.2% of the rat’s brain. A total of 14402 astrocytes populated the bounding region, co-localized with 88541 neurons and 1.37 m of vasculature wiring. The central microcircuit consisted of 15888 neurons, 2469 astrocytes, and 0.23 m of the vasculature, occupying a volume of 0.2 mm^3^.

### Data validation: Population-level

To ensure biological fidelity, we validated that input constraints could be reproduced for each step in the circuit building process and compared structural measurements with corresponding values extracted from the literature and experimental data. Astrocytes were placed according to the densities reported in Appaix et al. (2012), measuring an average density of 12241 mm^−3^, ranging from 9367 mm^−3^ to 21479 mm^−3^ close to the pia. The NGV framework accurately reproduced the density distribution for the selected profile (Figure 7A), corresponding to P14 rat data. Other studies reported similar numbers: 10700 ± 1750 mm^−3^ in African giant rats (Olude et al., 2015) and 10800 ± 400 mm^−3^ in mice (Schreiner et al., 2014). Astrocytic density increases from 2666 ± 133 mm^−3^ in neonates (Emsley & Macklis, 2006), to 15696 ± 860 mm^−3^ (Grosche et al., 2013) and 18000 ± 1750 mm^−3^ (Leahy et al., 2013) in adults. Densities in old rodents do not exhibit a significant increase with reported values of 18350 ± 1141 mm^−3^ (Nimmerjahn et al., 2004). Thus, the profile selection and generated densities lied within the range of juvenile numbers encountered in the literature (Figure 7E).

**Figure 7:**
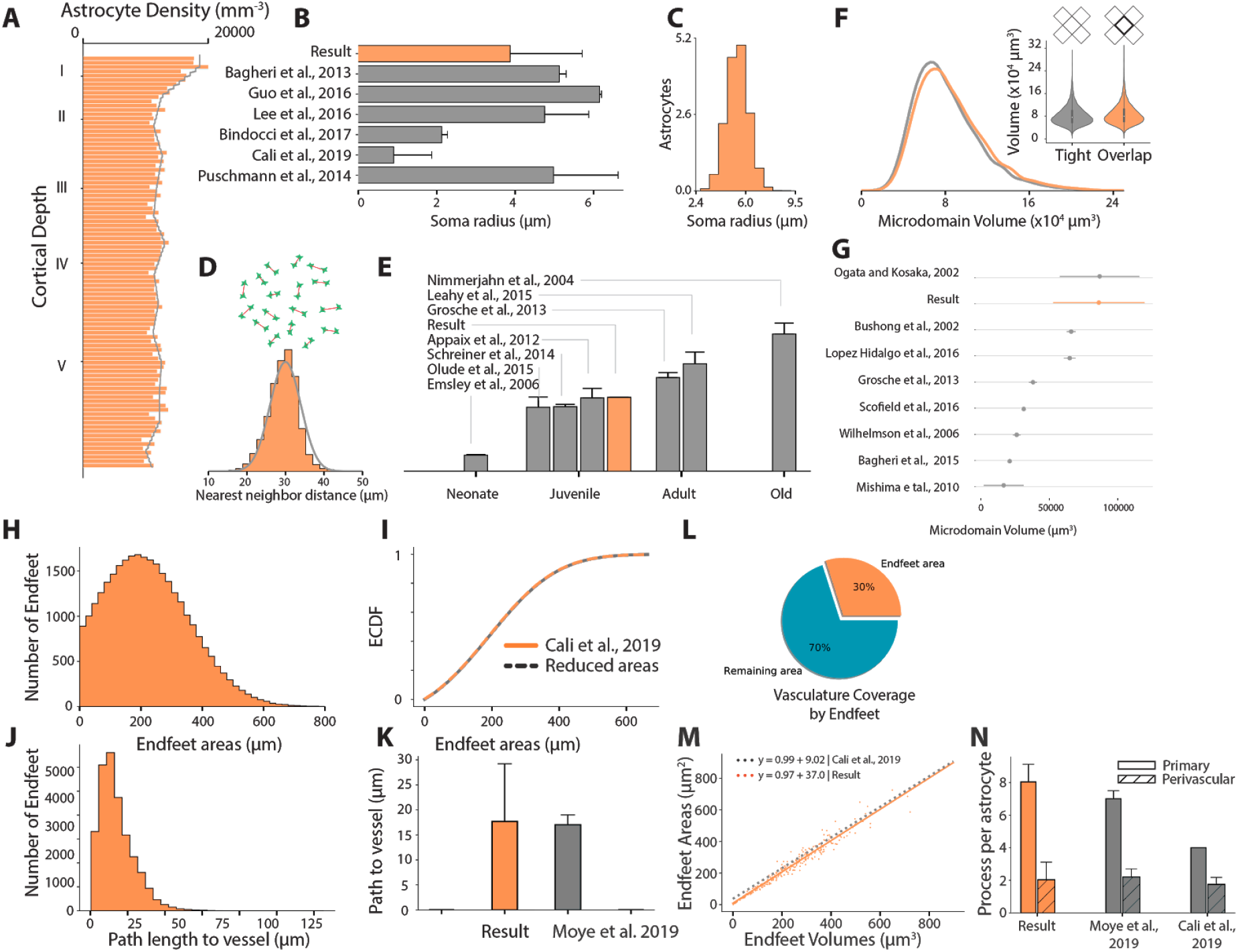
NGV Data Validation - Population-level. NGV circuit results are represented with orange, while literature data with gray until stated otherwise (A) Astrocytic somata density comparison between the NGV circuit and reported values from Appaix et al., (2012). (B) Bar plot comparison between the circuit’s soma radius distribution and literature measurements. (C) The respective histogram of somata radius distribution. (D) Circuit’s histogram of the nearest neighbor distance distribution compared to the input constraint of Lopez-Hidalgo et al. 2016 (gray line). (E) Average astrocyte density in context with measurements from various literature sources. (F) Comparison of volume distribution between tiling (gray) and overlapping (orange) microdomains. (G) Microdomain volume distribution comparison vs. literature sources. (H) Pruned endfeet areas histogram and validation of its ECDF versus the target distribution from Cali et. al., (2019). The distribution gives rise to a 30% coverage of the total vasculature surface area (L), leaving 70% uncovered (blue). (J) Shortest path length distribution from the soma to the vessel surface validated against the measurements in Moye et al., (2019). (M) Reproduction of the endfeet volumes and areas relationship as measured in Cali et al., (2019). (N) Comparison of the number of primary and perivascular processes with respect to the literature.

Next, we validated the spatial association among astroglial somata to verify their orderly distribution. The distance between each astrocytic soma and its closest neighbor was calculated and the resulting distribution reproduced the target distribution from López-Hidalgo et al., (2016), with an average nearest-neighbor distance of 30 μm(Figure 7D). During placement, the radii of the astrocytic somata were sampled from a normal distribution, which was fitted on the experimental values of astrocyte soma radii found in a multitude of studies (Bagheri et al., 2013; Bindocci et al., 2017; Calì et al., 2019; Puschmann et al., 2014)(Figure 7B,C). However, due to the small size of the datasets, we allowed for higher variance instead of using any single one of these values to constrain our sampling method. The final result of the placement is astrocytic somata with their size linked to their positioning, introducing two improvements over previous methods of placing the somata as points and then assigning the geometry (Keller et al., 2018). Soma size is taken into account while being placed so that intersections of somata with other somata and the vasculature are eliminated.

The microdomain tessellation, representing the anatomically-exclusive regions of the astrocytes, was generated from the position and size of the placed somata. The average microdomain volume was 81,725 μm^3^, ranging from 11,697 to 266,599 μm^3^, whereas the overlapping microdomains were measured at an average of 86,106 μm^3^, ranging from 12,324 μm^3^ to 280,890 μm^3^. The overlapping distribution exhibited a peak-to-peak shift of 2815 μm^3^ to the right, due to the higher domain volumes due to their overlap (Figure 7F). In the literature, ages seem to be a determining factor in the size of the microdomains. Studies using adult animals report volumes from 16,400 μm^3^ to 31,000 μm^3^ (Bagheri et al., 2013; Grosche et al., 2013; Mishima & Hirase, 2010; Scofield et al., 2016; Wilhelmsson et al., 2006). Juvenile rodent microdomain volumes have been observed to range between 65,900 μm^3^ and 86,700 μm^3^ (Bushong et al., 2002; Ogata & Kosaka, 2002). The NGV model produced domain volumes that corresponded to juvenile astrocytic densities, capturing the magnitude from the respective studies (Figure 7G). In the work of Lanjakornsiripan et al. (2018) specialized techniques were used to analyze astrocytic morphology, identifying a volume distribution ranging from 40,000 to 180,000 μm^3^. The most notable result of their quantification was that layer I astrocytes exhibited a smaller volume than in the rest of the layers (see supplementary material). The NGV circuit reproduced this observation, which resulted from the significantly higher densities in layer I. Our quantification showed that the average size of the domains decreases as the average astrocyte density increases. This result provides an invaluable insight: the contact-spacing organization of astrocytes in biology induces constraints of purely geometric nature. This allows for the abstraction of astrocytes into mathematical entities, i.e. tessellation regions, verifying our initial assumption that astrocytic domains can be modeled as such.

Before pruning, the endfeet surface meshes accounted for 91.1% of the vasculature surface. Studies using chemical fixation for their tissues reported a 70% − 100% coverage (Kacem et al., 1998; Korogod et al., 2015; Mathiisen et al., 2010; Simard et al., 2003) of the vasculature by perivascular endfeet. However, Korogod et al., (2015) showed that chemical fixation induces swelling of the astrocytic compartment, leading to increased coverage. They reported that coverage of the vasculature by astrocytic endfeet was 62.9 ± 1.5% using cryo-fixation, which is more likely to preserve the anatomical structures of the neocortex. An unconstrained simulation of growing the NGV endfeet surfaces produced a full coverage of the vasculature, however, in order to generate a biologically plausible distribution the areas were pruned with respect to the reported endfeet area distribution (Calì et al., 2019). After the pruning, the resulting endfeet mesh area distribution was 225 ± 132 μm^2^ in the range [0, 1000] μm^2^ (Figure 7H), matching the respective biological values (Figure 7I). The total area of the pruned meshes covered 30% of the total vasculature (Figure 7L), less than half the coverage reported by Korogod. This discrepancy could be due to the very few data points from which the endfeet area distribution was generated (n=7) assuming normality. The shortest path length from the soma to the surface of the vasculature was measured to be 17.23 μm on average, ranging from 0.41 μm to 110 μm (Figure 7J). The average path length is in agreement with reported values (Moye et al., 2019) (Figure 7J). Lastly, the relation between the surface area and thickness of the endfeet geometries validated that they were in agreement with the relationship from the study of Calì et al. (2019) (Figure 7M).

In the NGV circuit, the number of perivascular processes was constrained for juvenile rodents at 2 ± 1 processes. These numbers are in accordance with literature measurements (Calì et al., 2019; Moye et al., 2019). Furthermore, the number of primary processes in the NGV was measured as 8 ± 1 processes (Figure 7M). To validate the spatial relationship between neurons and astrocytes, for each astrocyte the distance to the closest neuronal soma was calculated 13 ± 5 μm, ranging from 0.7 μm to 30 μm. The distribution falls within the range of literature observations, that is from 5 μm to 30 μm for three types of inhibitory neurons (Refaeli et al., 2020).

### Data validation: Astrocytic Morphologies

Astrocytic morphologies were validated both on the single-cell and on the population basis to investigate the behavior of the synthesis algorithm in a vacuum, i.e. without the constraint of the microdomain geometry, and embedded in the circuit context-bound by the neighboring astrocytes. The validation was performed from both a morphometric/geometric and a topological point of view.

#### Clones of single astrocytes

Three clones were generated from each reconstructed astrocyte morphology, with each clone having as an input the original morphology’s endfeet appositions, morphometric distributions, point cloud, and branching topology (Figure 8A). As the synthesized cell was constrained by the original morphology, it allowed for the systematic study of its growth pattern without the external influence of the network organization. It should be noted that the synthesized cell morphology is still unique, regardless of the aforementioned constraints, due to the stochastic selection of the tree barcodes combined with the exponential probability of branching at a specific path length. For all the validations that follow, perisynaptic and perivascular trees were validated separately, due to their different roles and their distinct morphological properties.

**Figure 8:**
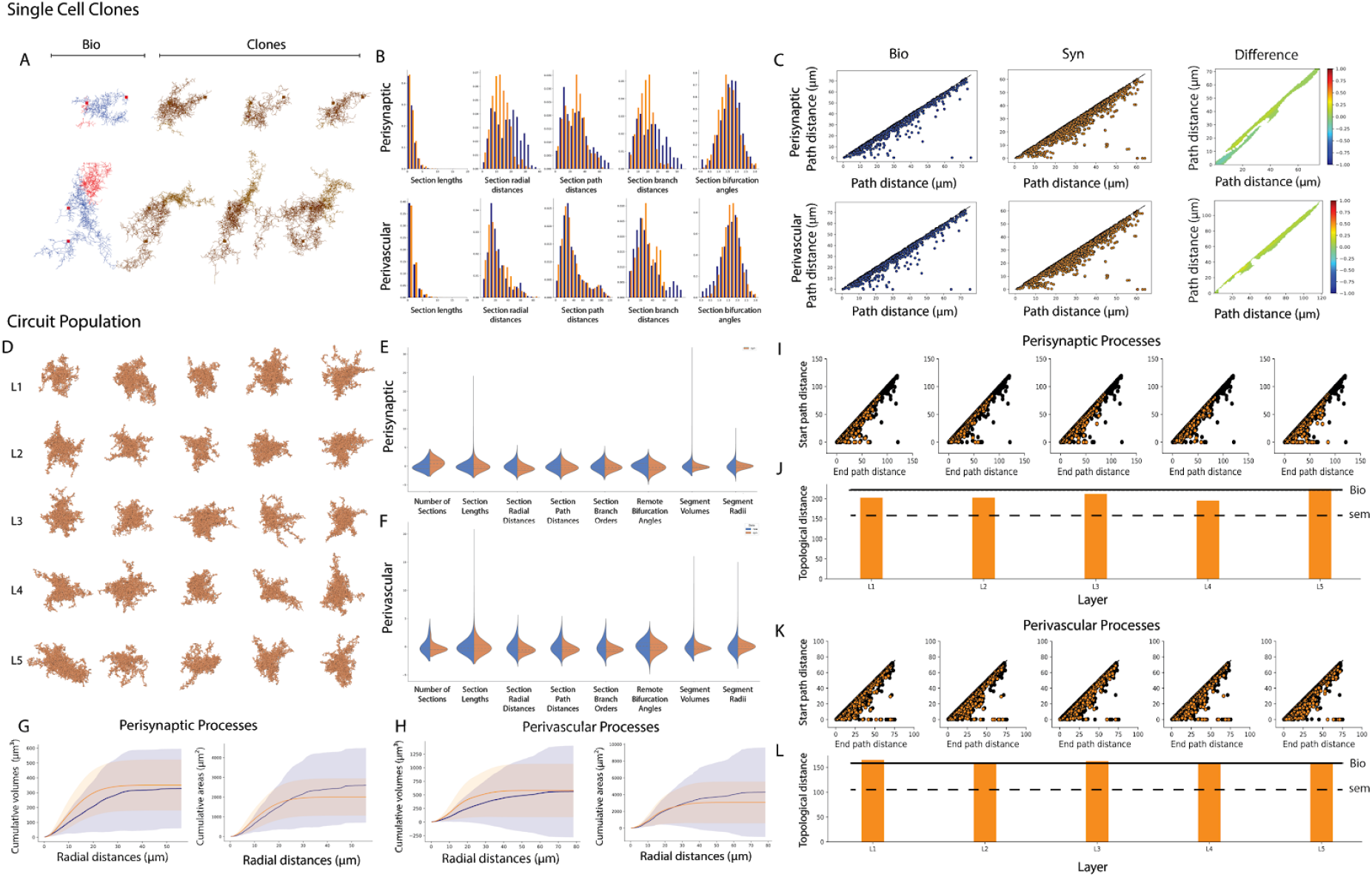
NGV Data Validation - Single-Cell Level Validation between reconstructed and synthesized morphologies. (A) Examples of synthesized cells (clones, orange) that are generated from a single branching topology, extracted from the digital reconstructions (bio, blue/red) on the left. (B) Comparison of morphometrics between clones (orange) and bio (blue) for perisynaptic (top) and perivascular astrocytic processes (bottom). (C) Persistence diagrams for bio and clones, and their topological difference for both process types. (D) Synthesized astrocytes examples from each layer from the circuit building framework. (E, F) Feature comparison between circuit astrocytes and bio for perisynaptic and perivascular processes. (I, K) Per layer persistence diagrams overlap between the synthesized and bio cells (J, L) The topological distance of each layer’s persistence diagram compared to the bio (black continuous line) and its respective standard error (black dashed line). (G, H) Validation of cumulative process surface area and volume as a function of the radial distance from the soma between circuit astrocytes (orange) and bio (blue).

A set of commonly used morphometrics was extracted from the two morphology groups. Specifically, we tested the section lengths, radial and path distances, branch orders, and bifurcation angles. When comparing morphometrics between groups of small size, as in this example in which the morphometrics of 3 cells are compared with the morphometrics of a single reconstruction, the mechanism of random barcode selection can result in creating the same tree topology more than once. This is especially likely if the pool of barcodes is created from a single cell. Therefore, the probability of drawing the exact combination of barcodes as in the reconstructed cell is very small. This effect can be seen in the morphometric distribution comparisons in Figure 8B as a mismatch in the distributions. Radial and path distances, as well as section branch orders, are the features that are most affected by the aforementioned bias. The effect is more prominent in the second cell, because of the larger difference between the barcodes of the trees. In all cases, the astrocyte successfully grew the perivascular processes to the endfeet targets using the available barcode. The topological colonization synthesis for astrocytes does not attempt to capture the morphometrics of a single morphology, which means to generate a morphology that is identical to the input cell. Instead, it aims to capture the population-wide characteristics while maintaining variability by creating unique realizations from that population. Therefore, the design choice of sampling from a pool of barcodes makes better sense when a population of cells is used as an input in synthesis as we will see in the next section. Before moving to a circuit wide validation, a more meaningful way to validate the correctness of the branching structure will be presented.

Persistence diagrams were extracted from the reconstructed and synthesized-cloned morphologies and subsequently converted to persistence images, the pixel values of which are normalized in the [0, 1] (Figure 8C). The size of the images in both axes is normalized by the maximum dimension (in μm) of our dataset. The topological difference between the branching topologies of the experimental and cloned cells showed a less than 20% discrepancy close to the diagonal of the persistence difference. Values close to the diagonal correspond to short-lived sections, i.e. their birth-death distance (section length) is small. This difference results from the choice of the segment length (0.1 μm), which leads to the scaling of all bars that are shorter than that threshold. Even though it contributes to a negligible increase of 0.62 ± 0.1 % of the total length of the cell, it is visible in the topological difference. The heavily ramified nature of astrocytic morphologies results in a high branching frequency, which requires a small segment length to capture all the details during synthesis. Having a ten times denser sampling of points makes synthesis significantly slower when compared to the neuronal synthesis that uses a segment length of 1 μm. However, it was necessary for capturing all the smallest bars. The dense morphologies can then be re-sampled after they are generated, reducing the number of points in longer sections and making sure that the short branches consist of at least two points, their beginning, and end.

#### NGV circuit synthesized astrocytes

In the context of a circuit, each astrocyte sampled its barcodes from all available branching topologies, which were previously extracted from the reconstructed datasets. The perivascular targets, perisynaptic orientations, synaptic clouds, and domain boundaries were generated from the NGV circuit building algorithms and were used as input in synthesis to guide the growth process. Therefore, the input data for each astrocyte was unique, determined by the local environment and the size of the astrocyte domain.

Five representatives from each layer that are not in contact with the boundaries were randomly selected (Figure 8D) from which a set of common morphometrics was extracted, such as the number and lengths of sections, the section radial and path distances and branch orders, the remote bifurcation angles and the segment radii and volumes. The same feature extraction was applied to the experimental reconstruction and the distributions of reconstructed and synthesized morphologies were compared for each feature.

Since the topological synthesis of astrocytes uses the path distance from the soma as a metric for branching, it was expected to reproduce very well the section length and path distance distribution for the population. Radial distance distribution, on the other hand, depends on the radial outgrowth of the synthesized trees. On the population level, the radial distances from all experimental morphologies were pooled together, smoothing out the distributions from longer or shorter cells. Therefore when compared with a synthesized sample that drew from barcodes randomly, similar statistics were achieved (Figure 8E,F). The simple rule of one child following the parent direction and the second child following the direction synapse that is available and furthest away from the initial direction was sufficient to reproduce the angle distributions of the experimental morphologies. This suggests that astrocytes create branches in a space-filling manner, as they progressively ramify during development, filling up the available space. Therefore, using the synaptic cloud, which is already available from the neuronal circuit, the biological splitting behavior can be reproduced without the need of extracting angle morphometrics from the experimental data, reducing in this way the input measurements that are required for astrocyte synthesis.

Simulation of calcium-induced waves throughout the morphology requires biologically realistic surface area and volume profiles for the distribution of ion channels and transmission line dynamics. For this reason, we computed how segment surface area and volume vary with respect to the radial distance from the soma of the astrocyte. First, we grouped all segments in each morphology in perisynaptic and perivascular regions and sorted each segment in the group according to the distance from the segment’s center to the center of the cell’s soma. The sorted-by-distance segment volumes and surface areas were cumulatively summed, producing the cumulative plots in Figure 8G. The synthesized diameters and perimeters resulted in a distribution of surface areas and volumes that fell within the variance of the reconstructed values. Note that the reconstructed cells assume significantly variable radial distributions, giving rise to high differences within the input population and rendering a proper modeling of diameters and perimeters difficult. More reconstructed datasets (10-20 astrocytes) would be necessary to successfully constrain the model and identify potential subtypes of astrocytes instead of pooling them together in the same group.

Similarly to the single-cell synthesized clones in the previous section, the trees of both reconstructed and synthesized morphologies were converted into persistence diagrams and subsequently into persistence images. The first step was to calculate the topological distance between each pair in the group of experimental morphologies in order to determine the error baseline within the reconstructed population. The average topological distance between reconstructed perivascular trees was 110 ± 32 units and for perisynaptic trees was 79 ± 24 units.

Next, we separated the synthesized cells into five groups, one for each layer, and we calculated the topological distance between each pair of experimental and synthesized morphologies and averaged them for each group. For perivascular processes, in layer I the average topological distance was 69 ± 27 units, in layer II 68 ± 33 units, in layer III 71 ± 40 units, in layer IV 65 ± 33, and in layer V 75 ± 33 units. For perisynaptic processes, in layer I the topological distance was 54 ± 11 units, in layer II 53 ± 9 units, in layer III 54 ± 11 units, in layer IV 52 ± 16 units, and in layer V 53 ± 12 units. These results can be seen in Figure 8I-L.

The comparison between the intra and per-layer inter-population topological distances showed that the reconstructed trees of the astrocytic morphologies exhibited large distances even when perisynaptic and perivascular processes were considered independently. One factor contributing to these results was the existence of trees in one morphology that were very short in radial extent as opposed to trees in a different morphology that were longer and spread out. Pruned trees could be the result of a partial reconstruction or cut from the reconstruction block. Given the small number of reconstructed morphologies (n=3), further classification or curation was impossible. However, the synthesized trees exhibited smaller topological distances on average because of the sampling of the barcodes.

### Data predictions

In experimental setups, there is a defined number of measurements that can be made, depending on the protocols that were used to stain, fix the tissue and digitally reconstruct it. In-silico anatomical reconstructions of the NGV do not seek to replace these types of experiments but to minimize the costs and time required for scientific discoveries. Algorithmically generated NGV circuits can serve as magnifying glasses into the brain’s complexity, allowing scientists to explore the geometry and topology of its cells and their connections. Moreover, the creation of multiple NGV circuits, each one with a different set of parameters that reflect organizational changes in brain anatomy, will allow for a better understanding of the anatomical principles and their geometric constraints. All these insights can enable the scientist to construct more focused experiments. Here we present an exploration of the quantification of some compositional and organizational aspects of the NGV circuit.

### Spatial organization of astrocytic endfeet

To gain a general overview of the spatial organization of the gliovascular elements, spatial kernel density estimate (kde) plots (Figure 9A) were generated from the points comprising these datasets. A Gaussian kernel was used for the estimation of the probability density function, the bandwidth of which (standard deviation) was determined by Scott’s rule (Scott, 2015). The plots were realized on the x-y plane, where y corresponded to the cortical depth of the circuit. The vasculature point samples were differentiated into large vessels and capillaries using a diameter threshold 6 μm (Schmid et al., 2019). In addition, two more datasets were used for the density plots: the astrocytic somata coordinates and the endfeet target points on the surface of the vasculature. The density plots showed no prominent spatial correlation between the endfeet targets and either the large vessels or the capillaries. They exhibited instead overlapping density regions with the density profile of the astrocytic somata, especially in layer I. In the NGV circuit most astrocytes produce endfeet (> 90% in the NGV circuit), therefore the higher density of endfeet in regions of high soma density was a consistent prediction. As a matter of fact the density of endfeet in the NGV circuit was measured as 23464 endfeet per mm^3^ in layer V and 28421 endfeet per mm^3^ in layer I. The cerebral microvasculature is a space-filling structure (Gould et al., 2011) that spans the entire cortex, whilst occupying less than 5% of the total cortical volume (Heinzer et al., 2006, 2008; Serduc et al., 2006). The astrocytic somata were evenly spaced following the density which almost doubled in layer one. Therefore, it appears that the generation of the endfeet was not restricted by the vascular volume and by extension most astrocytes always project endfeet to nearby vessels. Analyzing further the endfeet target point in layer I, we found that indeed the distribution of the endfeet targets wasn’t “trapped” by the vascular structures, rather it was homogeneously distributed throughout the available space. In addition, taking a closer look into a 30 μm slice verified that the endfeet target selection was spread out throughout the entire space, an observation that explained their spatial correlation with the astrocytic somata, but not with the vascular structures. In conclusion, the evenly spaced distribution of astrocytic somata throughout the neuropil allows for the generation of vascular endfeet projections, which extend to the vasculature from their local environment. The space-filling organization of the vasculature in combination with the astrocytic somata spacing allows for the uniform provision of glucose and nutrients to neurons (Magistretti & Allaman, 2018; Magistretti & Pellerin, 1996), which co-occupy the same space, and for an efficient recycling of water, neurotransmitters, waste molecules and ions (e.g. K + clearance) (Abbott et al., 2010; Bellot-Saez et al., 2017).

**Figure 9:**
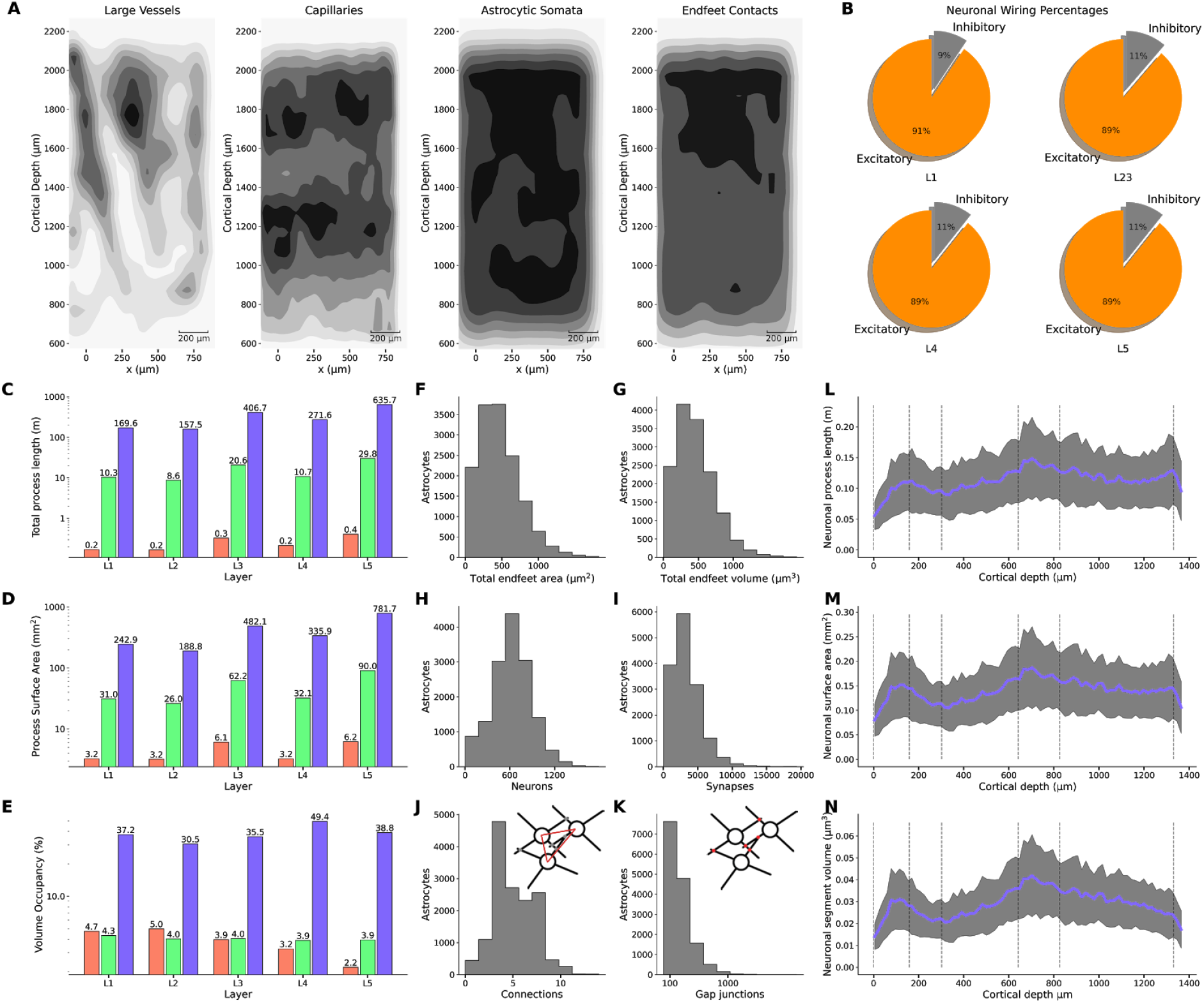
Data Prediction (A) Spatial densities of the points comprising large and small vessels, astrocytic somata centers and endfeet contact points. (B) Percentage of inhibitory (gray) and excitatory (orange) neurons per layer. (C) Neuron (blue), astrocyte (green) and vasculature (red) total process length, surface areas (D) and volume fractions (E) per layer. (F, G) total endfeet area dn volume per astrocyte. (H, I) Number of neurons and synapses connected per astrocyte. (J, K) Number of neighbors and gap junctional connections per astrocyte.(L-N) Total neuron process length, surface area and volume per astrocytic microdomain across the cortical depth.

### Astrocyte-related numbers

The reconstruction can predict quantities that are difficult to access experimentally. Due to the fact that the gliovascular interface has been extensively analyzed in the validation of the NGV circuit, two additional quantities were extracted that were not found in the literature, namely the total endfeet surface areas and volumes for each astrocyte. The median of the total endfeet surface areas was 427 μm (Figure 9F), and the median of the total endfeet volume areas per astrocyte was 414 μm^3^ (Figure 9G). The neuroglial interface consists of the connections between neurons and astrocytes via the formation of tripartite synapses. Each astrocyte domain connected to 627 ± 259 neurons (Figure 9H), whereas 6 ± 4 neuronal somata were in contact with each microdomain. The number of neuronal somata per astrocyte was in a consistent range compared to reported numbers of four to eight neuronal cell bodies per astrocytic domain in the rat hippocampus (Halassa, Fellin, Takano, et al., 2007). The median of the number of synapses per microdomain was 3010, which was less than the 100000 synapses per domain that have been reported by (Bushong et al., 2002). This discrepancy is due to the missing afferent fibers, mostly long range projections (Stepanyants et al., 2009). Taking into account the missing synapses from the external connections would result in a synapse density of 0.9 ± 0.1 μm^−3^ for layers I-V (Santuy et al., 2018), the predicted median for each microdomain would be 71995 and the 95th percentile 134010 synapses, which is consistent with the literature.

The astrocytic syncytium is formed via gap junction connections, established in the overlapping interface between each astrocyte and its neighboring astrocytes. The NGV circuit astrocytes formed 5 ± 2 connections with their neighbors (Figure 9I). The inter-astrocytic connections were reported as 11 ± 3 connections, ranging from 6 to 15 in hippocampal slices from P21 to P25 rats (Xu et al., 2010). In order to discern if the source of this difference in the emergent connections of the model was either a geometrical restriction resulting from the domain tessellation or morphological artifact, the number of neighbors per astrocyte was calculated based on the domain tessellation (using only the polygons). Therefore, analyzing the number of neighboring domains to each domain resulted in 15 ± 3 neighbors per astrocyte, which were notably higher than the connections established from the gap junctions. The connections were calculated from the detection of the intersections (touches) between neighboring morphologies. The median number of gap junctional connections were found to be 198, with the 95th percentile being 609 (Figure 9K). The exponential distribution of the gap junction numbers in combination with the available neighbors signified that the NGV astrocytes, not being yet fully mature, didn’t exhibit extensive ramifications, allowing for a uniform interface across the boundaries of the domain. The primary processes reached the boundaries of the domain and penetrated into neighboring territory forming clusters of connections. This is indeed how astrocytes form connections with their neighbors while developing and before reaching the maturation stage (Bushong et al., 2002, 2004; Ogata & Kosaka, 2002).

In silico circuits allow for a continuous and simultaneous analysis of multiple features. For example we estimated the average total wiring, surface area, and volume of neuronal branches per microdomain across the cortical depth (Fig. 10L-N). The total process density per microdomain was calculated for all neuronal morphologies and found to range from 1 to 2 μm / μm^3^ across all layers. The average number of neurons and synapses per astrocytic microdomain were 250 and 3000 respectively.

**Figure 10:**
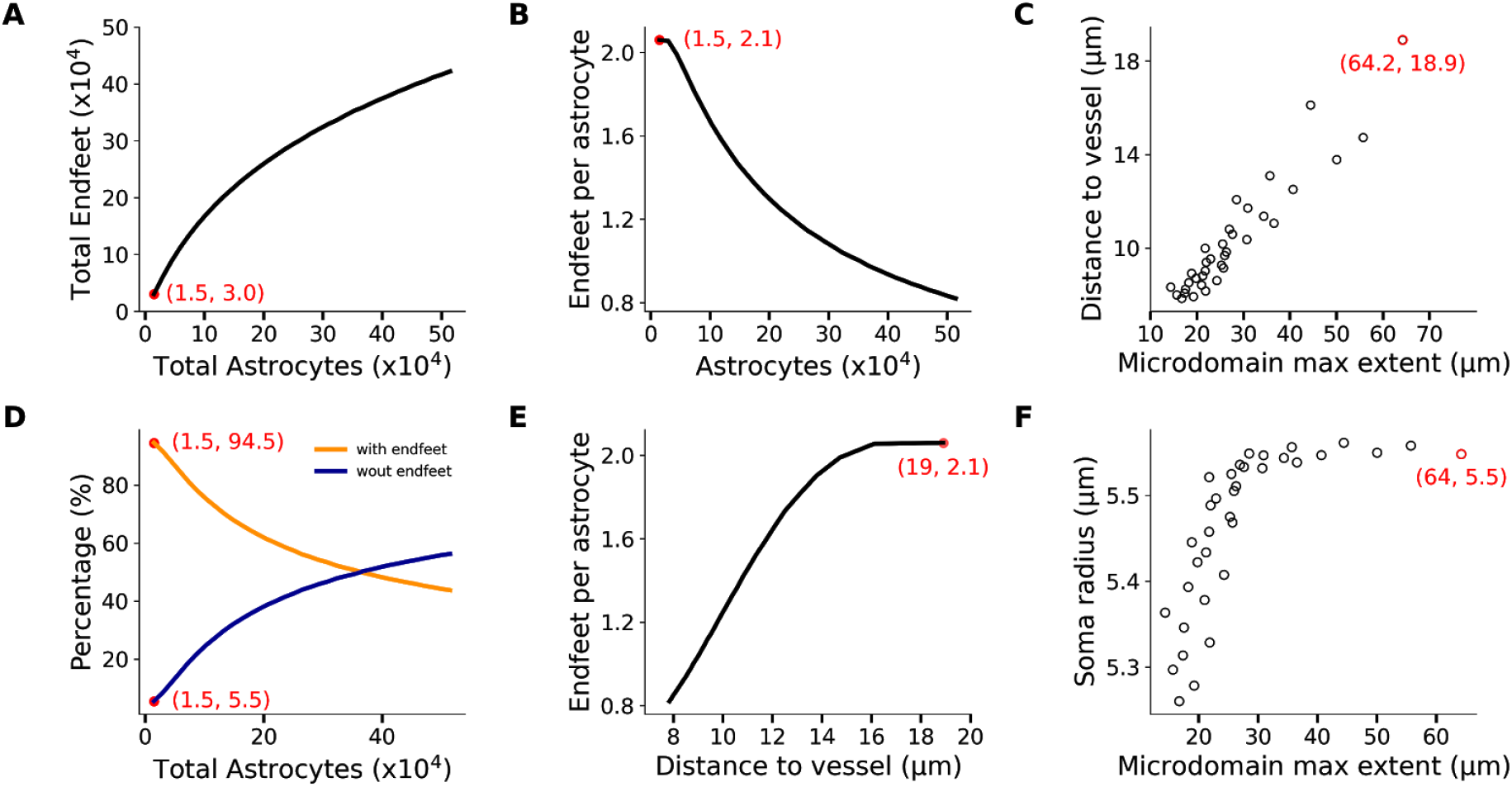
Effect of astrocyte density increase on the feasibility of perivascular processes in the same bounding space. The red data points correspond to the reference circuit with the biological parameters. As the astrocytic density increases the total number of endfeet increases sublinearly with respect to the total number of astrocytes (A), which is also reflected in the per astrocyte number of endfeet (B) and leads to smaller distances and domain extents (C). As the number of astrocytes increases, astrocytes with no endfeet increase in number (D), their distance to the closest vessel becomes smaller (E), and because of the packing, there is a bias for smaller soma sizes (F).

### Emerging NGV compositional hierarchy

In order to obtain a deeper understanding of the elements that comprise the gray matter, morphometric quantification was performed on the branches of neurons, astrocytes and the vasculature in the NGV circuit. More specifically the per layer total length, surface area and volume were calculated from the morphology segments (Figure 9C-E), along with their respective densities (see Table 1). The neuronal processes occupied 33 ± 13 %, astrocytes 4 ± 1 % and vasculature 3.8 ± 1.0 % of the neuropil volume. It has been reported that neuronal dendrites occupy 35 ± 5 % and axons 47 ± 5 % of the neuropil volume (Karbowski, 2015; Mishchenko et al., 2010). Astrocytes were reported to occupy 11 ± 4 % of the neuropil volume (Dienel & Rothman, 2020; Mishchenko et al., 2010), and vasculature less 5% of the total cortical volume (Heinzer et al., 2006, 2008; Serduc et al., 2006). The missing volume percentage of the neuronal wiring in the NGV circuit corresponds to the missing afferent fibers from outside the region of interest, which have been predicted to be approximately 41 million and would have formed an additional 147 ± 4 million synapses (Markram et al., 2015). Astrocytic process volume fractions exhibited a 6% less coverage compared to experimental estimates, which could not be explained in terms of missing wiring due to their localized structure. However, the NGV circuit models the anatomical architecture of the P14 rat somatosensory cortex, in which the astrocyte density (12286 ± 1601 mm^−1^) hasn’t yet reached adult values (15000 ± 18000, (Gordon et al., 2007; Leahy et al., 2013)). In addition, the degree of astrocytic ramification increases significantly from P14 to P21, at which age it converges into the mature spongiform phenotype that covers the entire domain (Bushong et al., 2002, 2004). The synthesized morphologies of astrocytes in the NGV circuit were generated from the branching topologies of P14 reconstructed astrocytes (Calì et al., 2019), which have not yet acquired the mature phenotype. Therefore, under the light of these two predicates, i.e. lower soma density and ramification compared to mature astrocytes, the lower volume fractions in the NGV circuit were reasonable compared to the reported measurements on adult rodents.

**Table 1.** Quantification of length, surface area and volume of the segments of neurons, astrocytes and the vasculature for each layer. Both total numbers and densities have been calculated.

| **Total length (m)** | **L1** | **L2** | **L3** | **L4** | **L5** |
| --- | --- | --- | --- | --- | --- |
| **Neurons** | 169.6 | 157.5 | 406.7 | 271.6 | 636.7 |
| **Astrocytes** | 10.3 | 8.6 | 20.6 | 10.7 | 29.8 |
| **Vasculature** | 0.2 | 0.2 | 0.3 | 0.2 | 0.4 |
| **Length density (m/mm<sup>3</sup>)** |  |  |  |  |  |
| **Neurons** | 1308.9 | 1347.7 | 1467.2 | 1824.4 | 1541.4 |
| **Astrocytes** | 79.2 | 73.7 | 74.2 | 71.5 | 72.3 |
| **Vasculature** | 1.3 | 1.4 | 1.2 | 1.4 | 0.9 |
| **Total surface area (mm<sup>2</sup>)** |  |  |  |  |  |
| **Neurons** | 242.9 | 188.8 | 482.1 | 335.9 | 781.7 |
| **Astrocytes** | 31.0 | 26.0 | 62.2 | 32.1 | 90.0 |
| **Vasculature** | 3.2 | 3.2 | 6.1 | 3.2 | 6.2 |
| **Area density (mm<sup>2</sup>/mm<sup>3</sup>)** |  |  |  |  |  |
| **Neurons** | 1874.9 | 1615.4 | 1739.4 | 2256.2 | 1895.4 |
| **Astrocytes** | 239.3 | 222.4 | 224.2 | 214.8 | 218.3 |
| **Vasculature** | 25.0 | 27.2 | 21.9 | 21.8 | 15.0 |
| **Total volume (mm<sup>3</sup>)** |  |  |  |  |  |
| **Neurons** | 4.8 | 3.6 | 9.8 | 7.4 | 16.0 |
| **Astrocytes** | 0.6 | 0.5 | 1.1 | 0.6 | 1.6 |
| **Vasculature** | 0.6 | 0.6 | 1.0 | 0.5 | 0.9 |

| Volume occupancy (%) |  |  |  |  |  |
| --- | --- | --- | --- | --- | --- |
| **Neurons** | 37.2 | 30.5 | 35.5 | 49.4 | 38.8 |
| **Astrocytes** | 4.3 | 4.0 | 4.0 | 3.9 | 3.9 |
| **Vasculature** | 4.7 | 5.0 | 3.9 | 3.2 | 2.2 |

Neuronal process total length ranged from 120 m in layer I to 657 m in layer V, two orders of magnitude higher than astrocytic process lengths, which ranged from 1.4 m to 3.5 m in layer V and three orders of magnitude higher than vasculature wiring, which ranged from 0.2 m in layer I to 0.5 m in layer V. Total surface area for neurons ranged from 242 mm^2^ in layer I to 781 mm^2^ in layer V. Astrocytic process surface area was measured from 31 mm^2^ in layer I to 90 mm^2^ in layer V. Finally, vasculature surface area ranged from 3.2 mm^2^ in layer I to 6.2 mm^2^ in layer V. The ratio of the total length between neurons and astrocytes was 20 ± 3 and between neuronal and vascular total length was 1347 ± 372. The total process surface ratios were 8 ± 1 and 98 ± 29, respectively. Following the quantification of the geometrical features of neurons, astrocytes and the vasculature, it was apparent that there was a systematic order-of-magnitude difference between them. The data suggested there is a hierarchy in cortical composition, the origin of which has been theorized in terms of length (Klyachko & Stevens, 2003; Wen et al., 2009), conduction delay (Budd et al., 2010), volume or/and spine economy (Karbowski, 2015) minimization. Most importantly, we showed here that an *in silico* circuit of the NGV architecture can indeed be used to investigate questions concerning the intricacies of cortical composition and their relation to computational capacity.

#### Effect of astrocytic density on endfeet organization

Many studies suggest that astrocytic density varies little across brain structures and species when compared to the variation of the neuronal density due to differences in neuronal sizes (Haug, 1987; Herculano-Houzel, 2014; Leuba & Garey, 1989; Tower & Young, 1973). As neurons increase in size in bigger brains, their density decreases leading to a higher glia/neuron ratio, ranging from 0.3 in rodents to 1.5 in humans (Blinkow & Glezer, 1968; Nedergaard et al., 2003; Pelvig et al., 2008). However, recent studies show that astrocytes in humans span twice the width as their rodent counterparts (Oberheim et al., 2009), which raises the question of how morphological constraints in a dense neuropil affect the astrocytic population.

In order to explore the structural dynamics between astrocytic density and microdomains, we generated multiple circuit realizations equipped with astrocytic somata, microdomains, and gliovascular connectivity, starting from a total of 14648 astrocytes and scaling up to 500 thousand, while keeping the bounding space, neuronal population and vasculature wiring unchanged. We found that as the astrocytic density increased, leading to a higher total number of astrocytes, the number of endfeet did not increase accordingly, but follows a sublinear relation (Figure 10A). This relation was a result of the reduction of the number of endfeet per astrocyte from an average of 2.1 to 0.8 (Figure 10B), induced by the shrinking of the microdomain bounding space, the extent of which dropped from 64.2 μm down to 15 μm (Figure 10C). A smaller microdomain extent reduces the reachable space of an astrocyte, which in turn results in the decrease of the number of astrocytes that have endfeet. Specifically, the percentage of astrocytes with no endfeet increased from 1.5% to almost 60% of the total astrocytes in the circuit (Figure 10D) and because of the tight packing, the average distance of the perivascular astrocytes to the closest vessel dropped from 19 μm to 0.8 μm (Figure 10E), with their somata essentially touching the surface of the vasculature and their anatomical domains occluding access to neighboring astrocytes. In fact, the packing becomes so dense that the Gaussian sampling of the somata radii becomes skewed, thereby favoring smaller values because of lack of available space (Figure 10F).

## Discussion

This study aimed to create for the first time a data-driven digital reconstruction of the NGV ensemble at a micrometer anatomical resolution that would allow for quantitative measurements and predictions, as well as the investigation of hypotheses concerning the structural architecture of the NGV. Using sparse data from numerous studies, we built an NGV circuit of 15888 neurons and 2469 protoplasmic astrocytes, forming a functional column of the P14 rat neocortex with its microvasculature.

The NGV construction consisted of five stages. First, we generated the positions and sizes of the astrocytic somata using biological distributions of densities, nearest-neighbor distances, and somatic radii. Next, we created a geometrical approximation of the astrocytic microdomains, establishing their tiling boundaries. From this tessellation, overlaps were introduced by uniformly scaling the domains until they reached a 5% overlap with their neighbors. The third stage included establishing the gliovascular and neuroglial connectivities using the microdomain geometry, which determined each astrocyte’s available search space. In the fourth stage, the endfeet vasculature sites were grown into surfaces that competitively wrapped around the vasculature’s surface, constrained by experimental area distributions. The final stage was to grow stochastic astroglial morphologies, constrained by the astrocytic data produced in the previous steps. To make this possible, we developed a novel algorithm that combined topological branching and space colonization using neuronal synapses as attraction seeds.

The generation of astrocyte morphologies using topology to reproduce their branching pattern is a paradigm shift from existing approaches (Savtchenko et al., 2018), limited by the number of experimentally-reconstructed morphologies. Instead, by extracting the branching topology from scarce data, the framework allows for an unlimited number of morphologies, which replicate the input population’s biological branching topology, yet grown into unique, space-embedded, and context-aware forms. In our dataset, cell co-occupancy is addressed on the somatic level, in which astrocytic somata do not intersect with the vascular dataset. Branches did intersect with each other because the neuronal circuit consisted of intersecting experimental reconstructions. To apply intersection avoidance strategies would require fully synthesized circuits (Torben-Nielsen & De Schutter, 2014; Vanherpe et al., 2016) in the future, in which all cells would grow simultaneously.

The NGv circuit was validated against numerous sources to ensure its biological fidelity. It successfully reproduced the spatial organization of the astrocytic somata, their overlapping volumetric domains as geometrical boundaries, the connectivity of the perivascular processes with the vasculature, and their respective endfeet areas and volumes. We found that the combination of constraints concerning the astrocyte densities, number of endfeet, and endfoot area distributions resulted in a total coverage of the vasculature equal to 30% of its entire surface. This indicated that this set of parameters could not give rise to the reported coverage of the vasculature, showing the synergistic constraints that may arise from a geometric architectural model.

For the validation of synthesized astrocytes, we extracted geometrical features and branching topology from experimental reconstructions and validated them against our synthesized population. We first investigated the morphological synthesis on a single cell basis, without the circuit context, validating the reproduction of the morphometrics and the persistence diagrams. The full circuit builds used all the available branching topologies from the reconstructions and the synthesized cells were embedded in the circuit context, with specific vasculature targets, synaptic cloud, and microdomain constraints. We validated that both morphometrics and topologies for perisynaptic and perivascular processes matched the biological distributions.

A fully-grown astroglial circuit, connected to both neurons and the vasculature, provided a window into the structural nature of astrocytes and how they can interact with neurons and the vasculature. The co-localization of neurons, astrocytes and the microvasculature allows for an integrated view of the NGV unit and provides a foundation for functional models of blood-brain-barrier (BBB) energetics and signaling, for calcium-induced waves throughout the astrocytic syncytium, modeling of blood flow, and for the establishment of tripartite synapses.

The NGV model provided a plethora of measurements that we collected as exploratory predictions of the underlying biological complexity. We found that the ubiquitous perivascular processes were not influenced by the vascular topology and that their distribution was only influenced by the proliferation of astrocytes. More specifically, the analysis of astrocyte somata’s spatial densities, vasculature, and endfeet unveiled that astrocytic endfeet are homogeneously distributed in space due to the space-filling geometry of the vasculature. This geometrically-constrained architecture allows for a spatially-continuous provision of trophic support to neurons throughout the cortical space, which only varies with cortical astrocytic density. Thus, the tiling astrocyte compartmentalizes the vasculature, and their endfeet optimize the communication wiring from the endfoot to neurons. Given the relatively low density of astrocytes compared to neurons, this endfeet organization and coverage would be impossible if astrocytes did not partition the cortical space with the anatomically exclusive regions, leading to insufficient trophic support.

Delving deeper into the NGV quantification, we extracted the per layer lengths, surface areas, and volumes, both in terms of total and density measurements. Compared to reported biological numbers, the volume fractions of neuronal processes were smaller than the values reported in the literature due to the missing afferent fibers that reached the circuit from outside. Also, the lower average density of the P14 rat neuropil combined with the partially ramified morphologies resulted in a volume occupancy that was 6% lower than reported values for adult animals. The quantification of the geometric features of all three elements in the NGV verified the emergence of a systematic order-of-magnitude difference in the cortical composition that has been observed in experimental studies. The NGV model renders possible the simultaneous quantification of both compositional (densities, wiring, surface areas, and volume) and organizational (connectivities, distance distributions, correlations) aspects of its entities.

Multiple circuit realizations of increasing astrocyte densities were generated to explore how endfeet appositions vary with respect to the number of astrocytes. Specifically, we evaluated the effect of increasing the astrocytic density, up to half a million astrocytes, on the network’s gliovascular connectivity. We found that as the total number of astrocytes increased, their overall extent shrank due to their tight packing, reducing their access to vascular sites. However, the astrocyte number increase did not compensate for the drop in endfeet numbers due to the tight packing of domains that prevented astrocytes from projecting to the vasculature. In contrast to neurons, astrocytic density varies little in different species and animal ages. Our experiments indicated that the contact spacing behavior, which gives rise to anatomically-exclusive domains, acts as a global constraint for the astrocytes’ morphological steady state, which is reached at one month of age in rodents (Bushong et al., 2004). In addition, for the morphological domain to include the vascular sites within reach, a specific range of spacing is required, which depends on the inter-vessel distance. Therefore, the astrocyte’s role in providing trophic support polarizes its morphology and constrains its location to maximize the connections from the vasculature to neurons.

Additional biological details will be incorporated as more data become available. This will improve the integration of diverse features and will impose additional constraints on the model, increasing its biological fidelity. Emerging studies have identified new subtypes (Clavreul et al., 2019, Batiuk et al., 2020) of protoplasmic astrocytes, the morphological characteristics of which may vary in different cortical depths and regions. Additional reconstructions of these new subtypes and their respective locations and densities are required in order to extract their topological characteristics and to improve the current recipe. Future iterations of the NGV architecture will expand into deeper regions, including fibrous astrocytes (Matyash & Kettenmann, 2010) and other types of macroglia, such as oligodendrocytes. Fibrous astrocytes grow straight and less branched processes, which connect to Ranvier’s nodes of myelinated axons and the vasculature much like their protoplasmic relatives (Marín-padilla, 1995). They cover larger regions (up to 300 μm) than protoplasmic astrocytes (< 60 μm) (Ransom, 2012) lack fine processes (Oberheim et al., 2009) and they are evenly distributed, although they don’t form anatomically exclusive regions (Vasile et al., 2017). During neuronal development, oligodendrocytes myelinate axons to increase signal conduction velocities and form bi-directional functional units (Nave & Trapp, 2008). One oligodendrocyte attaches to multiple axonal fibers and can be depolarized by them, influencing the axonal conduction velocity in a coordinating manner (Yamazaki et al., 2010), which is not yet understood. In addition, the inclusion of microglia (Davalos et al., 2005; Nimmerjahn et al., 2005), which play an important role in brain health and disease (Schafer et al., 2013), will introduce new computational challenges due to their constantly motile, dynamic nature.

Biophysical models of the BBB interface and its metabolic signaling require precise geometrical specifications of the astrocytic endfeet. Therefore, the algorithmic reconstruction of the endfoot-apposing surface allows for the modeling of the functional interface between astrocytes, pericytes, smooth muscle, and endothelial cells. The algorithmic approach of the present study allows for the reconstruction of all endfoot surfaces in a circuit, across the entire microvasculature in the region of interest. Changes in the endfeet surface areas, which lead by extension to the variation in the coverage of the vasculature, have been observed in pathologies such as major depressive disorder (Rajkowska et al., 2013) and Huntington’s disease (Hsiao et al., 2015). For example, the extent of the endfoot area determines total counts of the Kir and BK potassium channels that need to be distributed for a model of potassium buffering in the NGV unit (Witthoft et al., 2013). Therefore, a valid distribution of abutting endfeet areas is an integral part of a functional model of the blood brain barrier. A crucial factor for the accuracy of the endfeet areas is the quality of the vascular surface mesh. Disconnected components, floating segments, and reconstruction artifacts will all negatively influence the faithful reconstruction of the endfeet, as they will either trap the growth of an endfoot surface or force it to grow on a non-existent structure. In our model, vasculature reconstruction errors are present and influence the endfeet area distribution; however, with upcoming high-quality datasets, such sources of error will be reduced.

Our model provides the structural foundation for the large-scale biophysical modeling of cross-talk between neurons, glia, and the vasculature. This data-driven approach allows for incremental refinement as more experimental data become available, new biophysical models get published, and new questions arise. Drug delivery research studies the molecular properties of drugs, but should also take into account the interaction of the drug with its environment, i.e. the physicochemical properties as the drug travels through the BBB to various locations of a healthy and/or pathological brain. Similarly, research in neurodegenerative diseases such as Alzheimer’s disease target reactive astrocytes, the morphology of which is entirely transformed with variation in their ramification, overlap, and proliferation compared to healthy brains. Although local interactions can be studied, it is the emergent large-scale effects of changes in the lactate shuttle, glutamate recycling, synthesis of glutathione and overall disruption in homeostasis that can provide insights that will aid in the advance of therapeutic solutions. This model will provide a solid basis for this type of work.

## Acknowledgments

We thank Jay Coggan for the helpful conversations and for his editing contributions throughout the manuscript. We also thank Caitlin Monney for designing the table of contents illustration and cover artwork.

## Notes

**Conflict of interest statement:** No conflict of interest

**Funding:** This study was supported by funding to the Blue Brain Project, a research center of the École polytechnique fédérale de Lausanne (EPFL), from the Swiss government’s ETH Board of the Swiss Federal Institutes of Technology, and is based upon work supported by the King Abdullah University of Science and Technology (KAUST) Office of Sponsored Research (OSR) under Award No. OSR-2017-CRG6-3438.

### Competing Interest Statement

The authors have declared no competing interest.

