## supplementary for "Architecture of the Neuro-Glia-Vascular System"

### Supplementary Material

January 19, 2021

#### Astrocyte somata repulsion

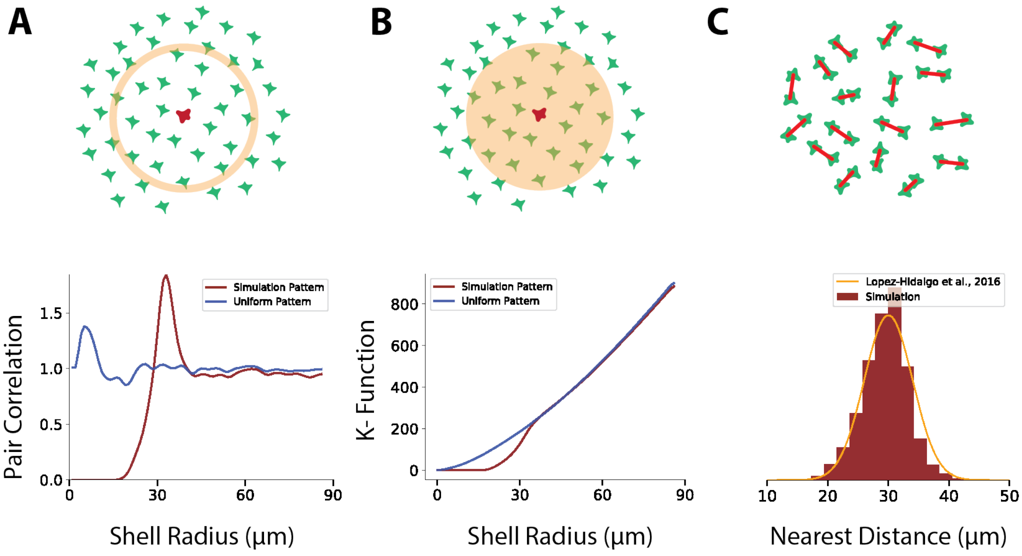


Figure 1 Spatial analysis of the point patterns corresponding to astrocytic somata. (A) Pair correlation function. (B) Ripley's K-function. (C) Distance distribution from each astrocytic soma to its closest neighbor, compared to the input profile (orange) from Lopez-Hidalgo et al. 2016.

Next, I validated the spatial association of the somata to verify whether they were evenly distributed. The simulation of contact spacing is essential for the formation of microdomains (Tout et al., 1993). I validated the repulsion model using three measures: pair correlation function (PCF), Ripley's K-function (Dixon, 2014) and nearest neighbor distribution. The PCF expresses the probability of finding one point a given distance from another point. Using the repulsive model, I found that the highest probability to find a soma at a distance around another soma was at 30 μm (Fig. 1A). In contrast, when the placement was ran without repulsion the respective PCF exhibited a small peak around 5 μm, the average radius of the astrocytic somata, rising from the algorithm's restriction for somata overlapping. Ripley's K-function is a similar spatial analysis method that uses a cumulative cover to describe the clustering or dispersion of a spatial point pattern. A random point sample is represented as a diagonal line, whereas clustered patterns move the line above the diagonal and disperse patterns move it below. Therefore, it was verified that astrocyte somata exhibit a disperse organization, while the non-repulsive placement demonstrated and almost diagonal trend, except from a small disperse deviation at very small distances because of the soma sizes. Finally, the nearest neighbor distribution explicitly quantified the distance of each soma to its closest neighbor, matching the input distribution at 30 μm.

Thus, all three measures verified that astrocytes were accurately spaced with respect to the target nearest neighbor distribution. Due to the fact that the repulsion functional can take any form, repulsive behavior is not the only possibility with this placement model. For example the spring and Lenard-Jones potentials have been successfully tested as well. In addition, more complex potentials that allow for repulsion and/or attraction to more than one types of elements (e.g. astrocytes and vasculature) can naturally extend the current model.

### Endfeet surface reconstruction & pruning

The growth of one endfoot was modelled as the solution $t(x)$ to the eikonal equation:

$$
\begin{aligned}
|\nabla_S t(\mathbf{x})| &= \frac{1}{f(\mathbf{x})},\ \forall \mathbf{x} \in \mathcal{S} \subset \mathbb{R}^3 \\
\phi(x) &= 0,\ \forall \mathbf{x} \in \partial S
\end{aligned} \tag{1}
$$

which is first-order partial differential equation, where S is the vasculature surface, a 2D smooth and closed manifold in $\mathbb{R}^3$, $\nabla_S$ is the gradient in the tangent plane to the manifold, $t(x)$ is the distance or travel time from the source and $f(\mathbf{x})$ is the speed of travel. Thus, $t(x)$ provides the time that an interface (contour) will need to reach $\mathbf{x}$ from the initial location $\partial S$. We are particularly interested in the simplified form where $f(\mathbf{x}) = 1$ and equation 1 is converted to the signed distance function from the boundary $\partial S$.

In our use case, in which we want to model the growth of an endfoot on the vasculature manifold, the boundary $\partial S$ corresponds to the endfoot target $\mathbf{x}_e$. Thus, the eikonal equation gives the travel times from the endfoot target to any point on S along the geodesics of its surface. Generalizing this to multiple endfeet targets, would required to calculate the travel times from each surface point to each endfoot target.

To approximate the solution of 1 on triangulation $S_T$ of the surface $S$, which is comprised of nodes $x_i$, I implemented the fast marching method for triangulated surfaces (Fu et al., 2011). Each node is assigned a value $T_i$ that corresponds to its travel time, which are initially set to $+\inf$ except for the endfoot's one which is 0. Using the one-ring neighbors for each node, the approximated solution of the travel $T_i$ at node $x_i$ is calculated from as the minimum of shortest path distances calculated from all the triangles in then neighborhood.For a triangle $(v_1, v_2, v_3)$, if $v_1$ and $v_2$ are upwind of $v_3$, there is a characteristic line of the gradient $\nabla_S t(\mathbf{x})$ that passes from $v_3$ and crosses the base of the triangle $\vec{e}_{1,2} = v_2 - v_1$ at $x_\lambda$. Thus, the travel time at $T_3$ is given by:

$$
\begin{aligned}
T_3 &= T_\lambda + T_{\lambda,3} \\
T_3 &= T_1 + \lambda\,(T_2 - T_1) + \|\vec{e}_{1,3} - \lambda \vec{e}_{1,2}\|
\end{aligned} \tag{2}
$$

which is derived from the fact that the approximation is linear, thus $T_\lambda = T_1 + \lambda T_{1,2}$ and $T_{\lambda,3} = f\|\vec{e}_{\lambda,3}\| = \|\vec{e}_{\lambda,3}\|$, because we have set the speed function to 1. To represent the gradient characteristic, $\lambda$ should minimize $T_3$ and must be in the range $[0, 1]$.

In order to introduce the notion of multiple endfeet growing in parallel, each node $x_i$ on $S_T$ was assigned an endfoot group $G_i$. Thus, upon initialization all endfoot nodes are assigned $T_i = 0$ and $G_i = i$, where $i = (0, 1, 2, ..., N_{endfeet})$. A priority queue was implemented that allowed to update first the nodes with the smallest travel time at each iteration, simulating the propagation of wavefronts. As nodes were updated with the shortest travel time to the nearest endfoot nodes, the group label of that endfoot was assigned to them and if the node had already a group assigned, the update stopped. This allowed the propagation of the "endfoot waves" on $S_T$ competitively as they were allows to propagate to nodes that were not already captured from a neighbor. The algorithm finished when there were no more vertices in the priority queue to update.

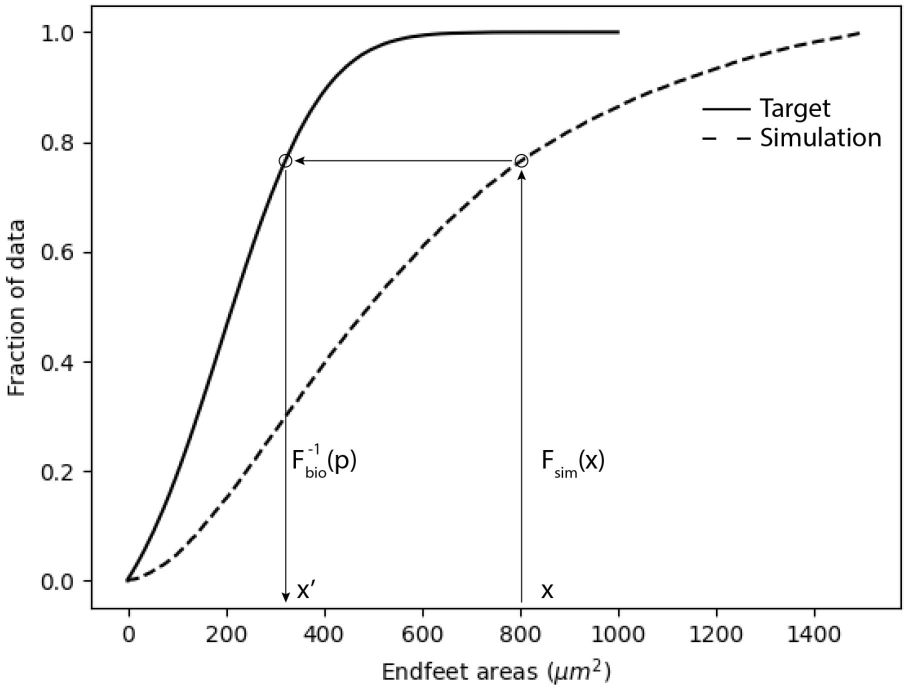


Figure 2 Diagram of the are transformation from the simulation area distribution to the target one extracted from the literature via the inverse CDF transform.

The endfeet meshes were reconstructed from the nodes in each group $G_i$. Due to the fact that the areas of the endfeet covered almost entirely the vasculature surface, a pruning procedure was introduced in order to match a target area distribution, extracted from the literature. Thus, given a target cumulative distribution $F_{bio}$ and the empirical distribution from the simulation $\hat{F}_{sim}$, overshoot endfeet areas $A_i$ were transformed into the respective target ones $A_i'$ via the cumulative inverse transform:

$$A_i' = F_{bio}^{-1}\left(\hat{F}_{sim}(A_i)\right) \tag{3}$$

To match the new pruned areas, the geometry of the endfoot meshes was pruned using the travel times that were calculated in the previous step. For each triangle in the endfoot mesh the average travel time was calculated from its vertices and the triangles with the highest travel times were removed one by one until the target area was reached. In other words, the meshes shrunk, starting from the periphery until the target area was approximately reached.

### Vasculature attraction field analysis

In order to model the chemo-attractive field which influences the growth of the perivascular processes, we need first to make some assumptions of its form and properties: We assume that there isn't a preferred direction of the diffusive gradient of the chemo-attractive molecules , i.e. it's isotropic. Furthermore, given that the vasculature graph and surface mesh are not available at the time of the morphology generation , the second assumption constrains the field to be generated by a point source instead of the entire surface of the vasculature in the vicinity.

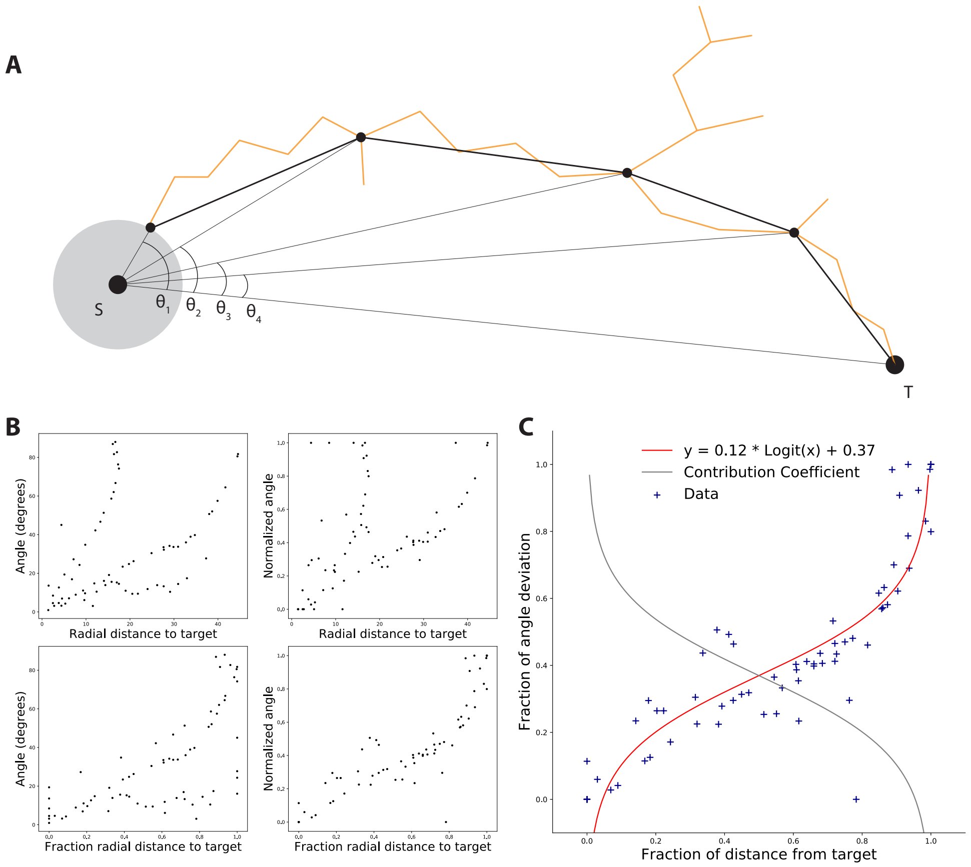


Figure 3 Attraction field analysis. (A) Analysis of radial distances from the soma S to each section end and the respective angle to $\vec{ST}$. (B) Scatter plots of non-normalized and min-max normalized data. (C) min-max normalized data (blue scatter) of how influenced the direction of the process is as it approaches the target along with the logit function fit (red) and the corresponding contribution coefficient $\alpha$

For each reconstructed astrocyte the endfeet targets were annotated as points close the the termination of an endfoot process. The closest leaf was found for each endfoot target and the upstream sections from the leaf to the root were extracted as show in Figure 3A.

Let a point $T$ be the target point, $S$ the center of the astrocytic soma. For each first point $p_i$ of each section, the angles $theta_i$ between $\vec{SP_i}$ and $ST$, as a function to the radial distance to the target $TP_i$. However, measurements on different trees lead to different trends that depend on the extent and orientation of the tree inside the attraction field. In order to normalize the data and quantify the underlying attraction trend we performed min-max normalization for both angles and radial distances as shown in the comparison of figure 3B.

The attraction of the main process to the target was fit using the quantile logit function (Figure 3C).

$$y(x) = 0.12 \times L(x) + 0.37 \qquad x \in [0, 1] \tag{4}$$

$$L(x) = \ln\left(\frac{x}{1-x}\right) \tag{5}$$

The function $a(x)$ produces values in the $(-\infty, \infty)$, but we are only interested in the interval $[0, 1]$. For this reason we introduce the clamp function $c(x) = \max(0, (\min(x, 1)))$, which limits the image into the desired interval. Finally, let $d_t$ and $d_{s_i}$ be the distances from the soma center to the target. The contribution factor of the direction to the target can be calculated, by substituting x with $1 - x$ and by using the identity $logit(1 - x) = -logit(x)$:

$$a\left(\frac{d_{s_i}}{d_t}\right) = c\left(y\left(1 - \frac{d_{s_i}}{d_t}\right)\right) = c\left(-0.12 \times L\left(\frac{d_{s_i}}{d_t}\right) + 0.37\right) \tag{6}$$

### Mean segment length estimation

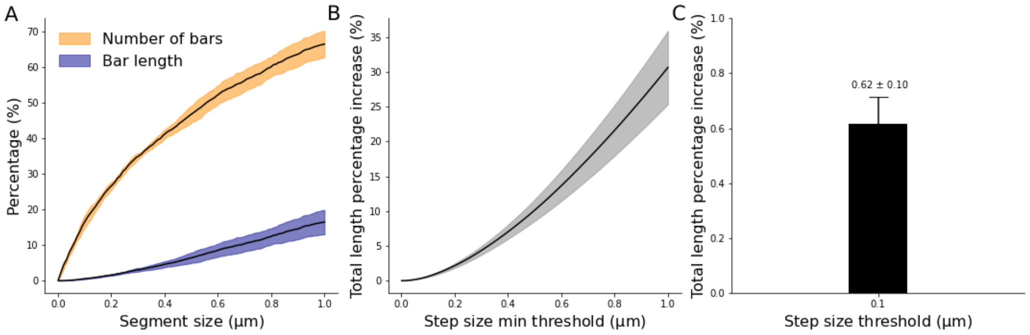


Figure 4 Bar length distributions. (A) Cumulative percentage of persistence bars that are smaller than segment size. (B) Total length increase in morphology with respect to minimum segment length and the exact value for segment length of 0.1 µm

In order to estimate the mean segment length, the cumulative percentage of the number of bars and bar lengths that are smaller than the segment size was plotted (Fig. 4A). Only a small fraction of bars (<10%) were smaller that 0.1 µm, indicating a good choice for the segment length threshold. The goal of this curation process isn't the removal of the smaller bars as it would remove branching points altering the topology, but rather the scaling of the bars below the threshold. The effect of the bar scaling was investigated in figure 4B, where the total length change of the morphology is plotted with respect to the chosen threshold and shows a negligible increase in total length which was quantified as $(0.6 \pm 0.1)$µm (Fig. 4C). A small $\sigma_{seg} = 0.001$ was chosen to allow for a small amount of variability.
